# Dimeric Transmembrane Orientations of APP/C99 Regulate γ-Secretase Processing Line Impacting Signaling and Oligomerization

**DOI:** 10.1101/2020.02.11.943480

**Authors:** Florian Perrin, Nicolas Papadopoulos, Rémi Opsomer, Devkee M. Vadukul, Céline Vrancx, Nathalie Pierrot, Steven O. Smith, Didier Vertommen, Pascal Kienlen-Campard, Stefan N. Constantinescu

## Abstract

Amyloid precursor protein (APP) cleavage by the β secretase produces the C99 transmembrane (TM) protein, which contains three dimerization-inducing Gly-x-x-x-Gly motifs. We demonstrate that dimeric C99 TM orientations regulate the precise cleavage lines by γ-secretase. Of all possible dimeric orientations imposed by a coiled coil to the C99 TM-cytosolic domain, the dimer (cc-del7) containing ^33^Gly-x-x-x-Gly^37^ in the interface promoted Aβ_42_ processing line and AICD (APP Intracellular Domain)-dependent gene transcription, including BACE1 mRNA induction, enhancing amyloidogenic signaling. Another orientation exhibiting ^25^Gly-x-x-x-Gly^29^ in the interface (cc-del6) favored processing to Aβ_43/40_, induced significantly less gene transcription, while promoting formation of SDS-resistant “Aβ-like” oligomers, reminiscent of Aβ peptide oligomers. These required both Val24 of a pro-β motif and the ^25^Gly-x-x-x-Gly^29^ interface. Thus, crossing angles imposed by dimeric orientations give access to γ-secretase at Aβ_48_ or Aβ_49,_ linking the former to enhanced signaling and Aβ_42_. We discuss avenues of blocking amyloidogenic processing.

## Introduction

The amyloid precursor protein (APP) is a type I transmembrane (TM) protein with a long extracellular domain, a single TM α-helix and a C-terminal cytosolic domain with no defined structure (Kang et al, 1987). Either α- or β-secretase first cleaves APP to generate two distinct fragments: a soluble N-terminal fragment (sAPPα or sAPPβ, respectively) and a membrane-anchored C-terminal fragment (α- or β-CTF, respectively) (Weidemann et al, 1989). Both α- or β-CTFs can be processed by γ-secretase to produce the APP intracellular domain (AICD), but transcriptionally active AICD is preferentially produced from the β-CTF or C99 (Belyaev et al., 2010) and only the β-CTF can generate the amyloid β (Aβ) peptides. These are present in various lengths ranging predominantly from Aβ34 to Aβ46. It is considered that Aβ40 is predominant in terms of production and that Aβ42 is the most toxic Aβ form, due to its higher propensity to aggregate because of the two extra hydrophobic residues at the C-terminus. Aβ42 forms the core of amyloid plaques in the brains of Alzheimer’s disease (AD) patients (Iwatsubo et al, 1994).

The first cleavage by γ-secretase on C99 occurs at the ε-site, corresponding to the residue 48 or 49 of the β-CTF, located at the cytoplasmic edge of the TM domain (Qi-Takahara et al, 2005; Takami et al, 2009). The ε-site is at a junction between the TM helix and the flexible cytosolic domain, where a helical to coil transition occurs at the structural level (Sato et al, 2009). Support for such an unraveling close to the ε-site and progressive cleavage was recently provided by the single particle electron microscopy structure of a complex between a C83 fragment cross-linked to catalytically inactive presenillin 1 (PS1) (Zhou et al, 2019). This leads to the formation of different AICD species, usually with cleavage occurring after residue 48 or after residue 49. The membrane-anchored fragment is further trimmed upstream by the γ-secretase carboxypeptidase activity. Two processing lines generate Aβ 46/43/40 and Aβ 45/42 fragments, respectively (Qi-Takahara et al, 2005; Takami et al, 2009). The AICD fragments have been described in gene regulation and require interaction with binding proteins in order to be translocated to the nucleus (Cao & Sudhof, 2001; Pardossi-Piquard et al, 2005). AICD signal transduction remains yet controversial. The way it binds to genomic sequences and its regulatory properties on transcription have been a matter of debate, but experimental evidence associates AICD release to the transcriptional regulation of APP target genes (Pardossi-Piquard et al, 2005).

APP contains three Gly-x-x-x-Gly motifs or glycine zippers at the junction between the juxtamembrane (JM) and TM regions. It is known from the glycophorin A protein that Gly-x-x-x-Gly and Gly-x-x-x-Ala particular motifs can induce SDS-resistant dimerization of TM proteins by creating close apposition between the glycines in the interacting α-helices (Bormann et al, 1989; MacKenzie et al, 1997). Importantly, SDS-resistant dimerization also requires other residues besides the Gly-x-x-x-Gly motif, and the position of the motif relative to the thickness of the bilayer is also important (Bormann et al, 1989; MacKenzie et al, 1997). The Gly-x-x-x-Gly motifs were reported as key molecular determinants that stabilize the association of the α-helices of APP TM dimers, and that modulate APP processing by the γ-secretase complex (Kienlen-Campard et al, 2008; Munter et al, 2007). The Flemish mutation in which there is an Ala to Gly substitution at position 21, creates a fourth upstream in-register Gly-x-x-x-Gly motif and induces a conformational change by increasing the α-helix at position 21 (Tang et al, 2014). This promotes an increase in Aβ production, confirming the relation existing between APP TM dimerization and APP amyloidogenic processing. Still, the precise mechanisms by which TM dimerization regulate the processing (and likely the function) of APP are not understood.

Here we asked whether dimers of C99 could be processed by γ-secretase and whether the precise dimeric interface of C99 TM helices could influence processing by γ-secretase leading to AICD formation and signaling via ε-cleavage, as well as Aβ accumulation subsequent to processing at γ-sites. We used fusions with a coiled coil dimer (Put3) at different junction points within the TM domain. These Put3 dimeric constructs allowed us to individually dial-in all seven possible dimeric interfaces in the C99 TM domain. The effects of each of the fusion proteins, reproducing one at a time the individual dimeric interfaces, were tested in cell lines and primary cells with respect to signaling, processing and oligomerization. We identified a precise dimeric interface where strong AICD release by γ-secretase processing leads to signaling and likely favors the Aβ42 processing line. Another interface rotated clockwise by 110° is associated with oligomerization of processed amyloid peptides and favors the Aβ43 processing line. Strikingly, both of these interfaces are lined with glycines and the shift from signaling to oligomerization involves the displacement of the crossing-point of the two helices from the third Gly-x-x-x-Gly motif in the middle of the TM domain, to further up towards the first Gly-x-x-x-Gly motif at the extracellular juxtamembrane-TM domain border. This difference in TM packing appears to be translated into processing by γ-secretase at different exposed sites and different oligomerization properties. We discuss how this knowledge can be translated in therapies aimed at preventing amyloid deposition, and we reveal that signaling and processing/oligomerization are supported by close dimeric interface.

## Results

### Design of the fusion proteins Put3 coiled coil C99

The left-handed leucine zipper coiled coil of the transcription factor Put3 (Put3-cc) from Saccharomyces cerevisiae was used to create fusion proteins at different junction points within the extracellular JM domain of C99. This strategy has previously been used to determine the effects of dimeric orientation on type II TM proteins (Mattoon et al, 2001), where the N-terminus is cytosolic, and on type I TM proteins where the N-terminus is extracellular (Seubert et al, 2003; Staerk et al, 2011). For the erythropoietin and thrombopoietin receptors, it has been shown by cysteine-mediated cross-linking that dimerization at different orientations imposed at the extracellular end of the TM domains is translated into the correct interfaces at the C-terminal ends of the TM domains (Staerk et al, 2011).

We replaced amino acids 1-17 of the C99 extracellular domain with the Put3-cc, (Walters et al, 1997), which exhibits an overall neutral charge that makes it compatible with an extracellular topology. We engineered fusion proteins where the extracellular Put3-cc forces the C99 monomers to adopt each of the seven possible symmetric left-handed orientations downstream of the coiled coil, denoted cc-del0 to cc-del6 (cc-C99^18–99^ to cc-C99^24–99^ respectively). We also created three shorter constructs, denoted cc-del7 to cc-del9 (cc-C99^25–99^ to cc-C99^27–99^ respectively), where the interfaces are similar to the first three fusion proteins, but where significant residues of the JM region are deleted, as certain motifs such as ^16^LVFF^20^ have been reported to exert an inhibitory effect on processing by γ-secretase (Tian et al, 2010) (Figures 1A and 1B). All cc-C99 (cc-del(X)) fusion proteins and wild type C99 are HA-tagged and contain a signal peptide at the N-terminus. The position of heptad repeats characteristic of left-handed coiled coils are denoted a-g with interface residues in “a” and “d” positions, with edge residues close to the interface in “e” and “g” positions and with residues outside of the interface in “b”, “c” and “f” positions (Figure 1C). The cc-del0, cc-del1 and cc-del2 constructs exhibit the same orientations as the cc-del7, cc-del8 and cc-del9 constructs, respectively.

**Figure 1.**
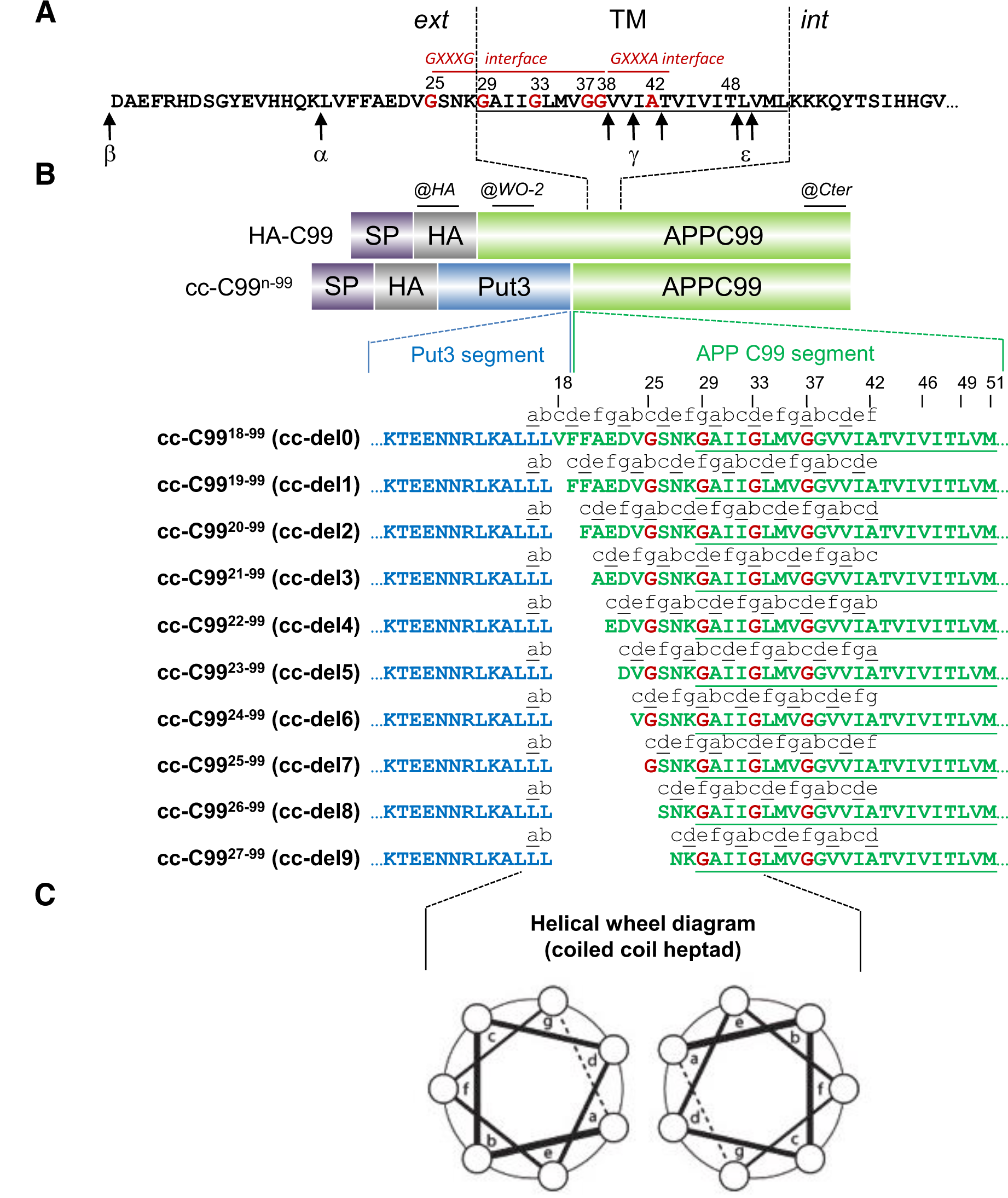
Design of coiled coil APP C99 Fusion Proteins. **A.** Representation of the juxtamembrane and transmembrane domains of human APP C99. The positions of the consecutive GxxxG motifs in Aβ numbering are in red. The cleavage sites of α-, β- and γ-(γ and ε) secretases are indicated by arrows. Ext: external, Int: internal, TM: transmembrane. **B.** Representation of HA-C99 and cc-del(X) constructs. The coiled-coil dimerization domain of Put3 was fused to consecutive residues of the juxtamembrane sequence of C99 starting at residue 17 (Aβ numbering) and where n is the first amino acid of the sequence. All seven possible registers of C99 TM dimers were imposed by successive N-terminal deletion. Constructs cc-C99^18–99^ to cc-C99^27–99^ are referred to as cc-del(X) with X from 0 to 9 respectively. cc-del7 to 9 have the same orientation as cc-del0 to 2 respectively, but without the ^16^LVFF^20^ motif. All fusion proteins contain a signal peptide (SP) and a HA tag upstream the Put3 sequence. Only HA-C99 harbors the WO-2 epitope. The Y188 antibody (Cter) is directed against the C-terminus of the C99 sequence. **C.** Helical wheel diagram showing the heptad repeats in cc-del(X). Position of heptad repeats are denoted a to g with the dimeric interface considered to be at positions a and d.

### Validation of the TM orientations of fusion proteins cc-del(X)

The dimer interface can be predicted for each fusion protein (Supplementary Figure 1A and 1B). To test the model of the predicted orientation of the cc-del(X) constructs, a cysteine was substituted for Gly29 (G29C) in all the constructs (Supplementary Figure 1A and 1B). Because this Cys residue is unique in the cc-del(X) fusion proteins, its position can be used as a read out for the dimerization interface by cross-linking studies. We used the Cys-dependent and irreversible cross-linking agent N, N’-1,2-phenylenedimaleimide (o-PDM) that forms dimers if two proteins are separated by a distance of 6 to 10 Å. We have predicted, based on *α*-helical wheel diagrams, which constructs can form dimers due to the presence of Cys residues within the interface itself (represented in red), with Cys residues in close enough proximity (represented in blue) or with Cys residues too far apart to dimerize (represented in green) (Supplementary Figure 1B).

Cross-linking experiments were performed with living cells and validated the orientations imposed by the coiled coil structure in the JM domain. Western blot analysis using the C-terminal (Cter) APP antibody showed all cc-del(X) G29C constructs were expressed in CHO cells at the expected size of 14 kDa (Supplementary Figure 1C). Cells treated with o-PDM are able to form intermolecular bonds maintaining stable dimers in denaturing and reducing conditions. In the presence of o-PDM, we observed the formation of cross-linked dimers for cc-del3 G29C and cc-del6 G29C, consistent with the hypothesis that these two constructs should have G29C in the dimeric interface (positions “d” and “a” of a heptad repeat). The cc-del0, cc-del2, cc-del7 and cc-del9 were also able to form dimers, the two cysteines being close enough to the interface via edge residues, placed at “e” or “g” positions of the heptad repeat. Specifically, G29C will be located in “g”, “e”, “g” and “e” for cc-del0, cc-del2, cc-del7 and cc-del9, respectively. The other dimers that do not crosslink exhibit C29 at positions “f”, “c” and “b”, namely cc-del1, cc-del4, cc-del5 and cc-del8, respectively. To validate that our constructs obeyed the predictions downstream of Gly29, we introduced a Cys mutation at position 37 (G37C) in cc-del4, cc-del6, cc-del7 and cc-del9. In the presence of o-PDM, we observed dimers in cc-del4 G37C and cc-del7 G37C but not in cc-del6 G37C and cc-del9 G37C (Supplementary Figure 1C). This is in agreement with our predictions as G37C was expected to be in the dimeric interface for cc-del4 and cc-del7 (positions “d” and “a” of the heptad repeat, respectively) but outside the interface for cc-del6 and cc-del9 (position “b” and “f”, respectively).

While the TM domain of C99 adopts an *α*-helix secondary structure (Nadezhdin et al, 2012; Tang et al, 2014; Zhou et al, 2019), it has been demonstrated that unraveling occurs upstream of the ε-site, starting around residues 48-49 (Sato et al, 2009). This unraveling is followed by a short cytosolic β-strand downstream, which is induced by the contact with PS1 in the γ-secretase complex (Zhou et al, 2019). We asked whether our engineered dimers allow this level of flexibility or whether they are maintained in a rigid conformation at the C-terminus of the TM domain. For this purpose, we substituted L49 with cysteine (L49C) in all ten cc-del(X) fusion proteins. Cross-linking with o-PDM could be detected for all fusion proteins, consistent with unraveling at that position, similar to that described for wild-type C99 (Supplementary Figure 1D) (Sato et al, 2009; Zhou et al, 2019).

### cc-del7 favors transcriptional activity via AICD production by γ-secretase activity

The release of transcriptionally active AICD was measured in CHO cells which were co-transfected with the HA-C99 or cc-del(X) fusion proteins, Fe65, Tip60Gal4, Gal4RE-luciferase as a reporter gene (Cao & Sudhof, 2001) and the pRLTK (renilla luciferase) control reporter to normalize for transfection efficiency (Figure 2A). The level of transcription was significantly increased as expected with the HA-C99 when compared to the mock control, but strikingly only with cc-del7 among the cc-del(X) constructs (Figure 2B). For the other constructs, the level of transcriptional activity was not significantly different compared to the mock control. It appears therefore that only one precise TM dimeric orientation (cc-del7) promoted APP-dependent nuclear signaling.

**Figure 2.**
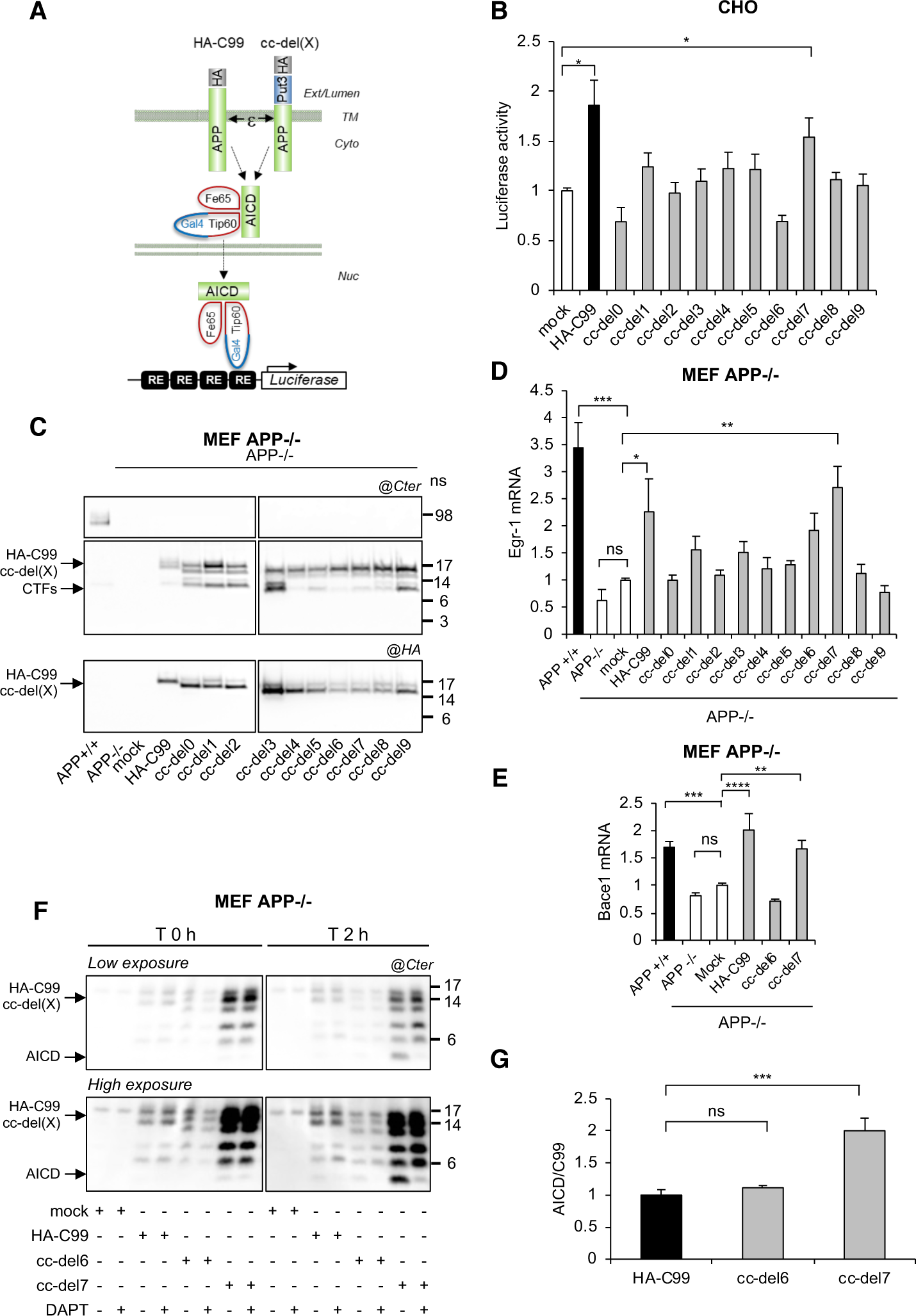
Signaling and gene expression via AICD production of cc-del7 fusion proteins. **A.** Transactivation assay to monitor the release of transcriptionally active AICD. Cleavage at the ε-site releases AICD, which translocates to the nucleus together with Fe65 and Tip60Gal4. The AICD/Fe65/Tip60Gal4 complex transactivates the Gal4RE reporter gene. Ext: exernal, Cyto: cytoplasm, TM: transmembrane, Nuc: nucleus. **B.** Transcriptional activity of the cc-del(X). Data are mean values ±SEM (n = 3, N = 3). Statistical analysis: non-parametric multiple comparisons Steel’s test with control (mock). *p<0.05. **C.** Western blots of HA-C99 and cc-del(X) constructs stably expressed in MEF APP-/-, revealed with the APP Cter (top) or the HA antibody (bottom). **D.** *Egr-1* mRNA of MEF cells described above (C). Results (Means ±SEM; n=3; N=3) are given as fold induction compared to control (APP-/-mock). **E.** *Bace1* mRNA of MEF cells described above (C). Results (Means ±SEM; n=3; N=3) are given as fold induction compared to control (APP-/-mock). **F.** AICD production in MEF APP-/-expressing cc-del(X) constructs. Total membrane extracts analyzed by Western blotting and revealed with APP Cter. **G.** Quantification of AICD. Results (Means ±SEM; n=3; N=2) are given as AICD:HA-C99 or AICD:cc-del(X) ratios, with AICD:HA-C99 ratio fixed at 1. Statistical analysis (**D, E** and **G**): one-way ANOVA followed by Dunnett’s multiple comparison test. *ns*: non-significant, *: p<0.05, **: p<0.005, ***: p<0.001, ****: p<0.0001.

To determine whether this effect was truly due to the activity of cc-del(X) and the release of transcriptionally active AICD, we created stable cell lines using murine embryonic fibroblasts deficient for APP (MEF APP-/-). Western blotting with antibodies directed against the C-terminus of C99 (Cter) or the HA epitope indicated that cc-del(X) fusion proteins exhibited comparable levels of expression in MEF APP-/-cells (Figure 2C).

Within these stable cell lines, qPCR was used to analyze the expression of the *Erg-1* gene known to be an APP downstream target (Hendrickx et al, 2013). *Egr-1* is an immediate early response gene regulating the transcription of late response genes important for synaptic plasticity processes, especially the maintenance of long-term potentiation (Jones et al, 2001). Exposure of mice to a novel environment induces *Egr-1* expression in an APP-dependent manner (Hendrickx et al, 2013). We therefore chose *Egr-1* as it is transiently induced via APP, and appears as a readout for the signaling function of APP upon cleavage by γ-secretase. We observed a decrease in the expression of *Erg-1* in MEF APP-/-compared to MEF APP+/+ cells (Figure 2D). The expression of *Erg-1* was restored in MEF APP-/-cells expressing HA-C99 or cc-del7, but not the other constructs. Importantly, Egr-1 is a transcriptional activator of the ß-secretase (*Bace1*) and its inhibition decreases amyloidogenic processing in Alzheimer mouse models (Qin et al, 2016). We used qPCR to assess whether *Bace1* expression was also increased in cells stably expressing cc-del7. The expression of *Bace1* was decreased in MEF APP-/-compared to MEF APP +/+. Its expression was restored in MEF APP -/- cells stably expressing HA-C99 or cc-del7 but not in MEF APP-/- cells stably expressing a mock vector or cc-del6 (Figure 2E). In agreement with previous results (Qin et al., 2016), the expression profile of *Bace1* mimicked the one observed for *Egr-1*.

Knowing that APP-dependent gene expression can be mediated by transcriptionally active AICD, we next assessed the production of AICD via a cell free experimental approach. Whole cell membranes from MEF APP-/- cells expressing different cc-del(X) constructs were isolated with hypotonic buffer and ultracentrifugation. The membrane lysates were analyzed by Western blot using an antibody directed against APP C-terminus at two time points, 0 h and 2 h of incubation at 37 °C in the absence or presence of a γ-secretase inhibitor (DAPT). The results indicated an increase in intensity of the band corresponding to AICD after 2 h compared to the initial time-point for all constructs tested. Strikingly, this increase was much stronger for cc-del7 than for cc-del6 or HA-C99 (Figure 2F). Quantification of the ratio of AICD produced confirmed a significant increase for cc-del7 when compared to the other constructs (Figure 2G). The bands considered (below 6 kDa) were strongly reduced in the presence of DAPT, indicating that they truly correspond to AICD produced by the γ-secretase .

### cc-del7 orientation favors AICD-dependent signaling in human neuronal cell lines and in rat primary neurons

To test whether the cc-del(X) fusion proteins exerted the same effects in neuronal cells, we used the SH-SY5Y human neuroblastoma cell line, which exhibit neuronal properties upon differentiation with retinoic (Agholme et al, 2010). We generated SH-SY5Y cell lines stably expressing cc-del(X) by lentivral transduction followed by puromycin selection. The expression profile of fusion proteins was tested 7 days after differentiation by Western blotting (Figure 3A).

**Figure 3.**
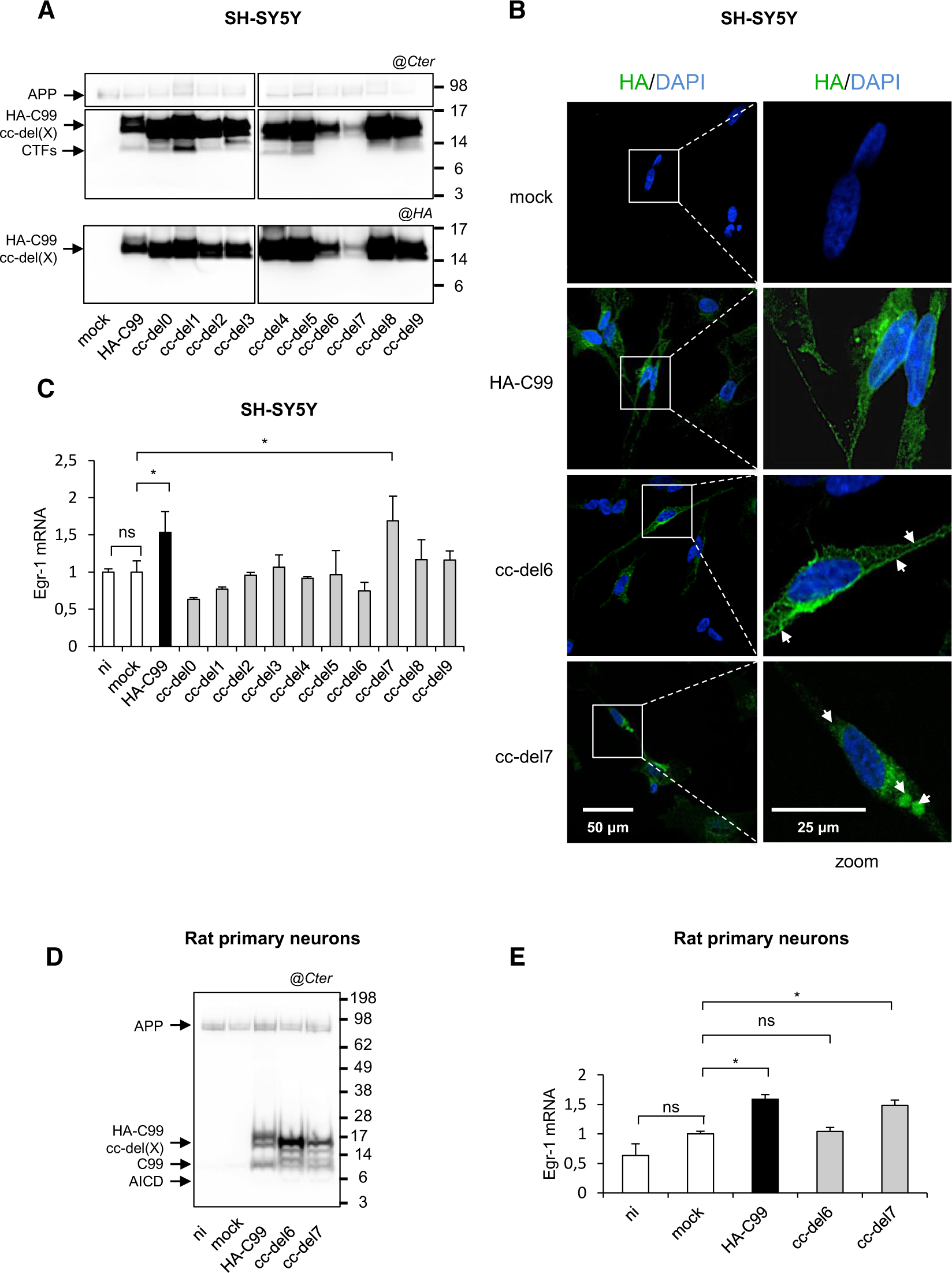
Regulation of the transcriptional activity by the cc-del7 construct. **A.** Cell lysates SH-SY5Y cells stably expressing HA-C99 or cc-del(X) constructs analyzed by Western blotting and revealed with the APP Cter antibody (top) or the HA antibody (bottom). **B.** Cells were differentiated by retinoic acid treatment for 7 days, stained with HA antibody (green) and DAPI (blue) and analyzed by confocal microscopy. White arrows indicated localization of cc-del(X) constructs. Left scale bars: 200 μm, right scale bars: 100 μm. **C.** *Egr-1* mRNA levels in SH-SY5Ycells. Results (Means ± SEM; n=3; N=3) are given as fold induction compared to control (mock). Statistical analysis: one-way ANOVA followed by Dunnett’s multiple comparison test. *p<0.05. **D.** Rat cortical neurons were transduced with lentivirus expressing mock vector (GFP), HA-C99, cc-del6 and cc-del7 or non-infected (ni). Neuronal lysates were analyzed 4 days after infection by Western blotting and revealed APP Cter antibody (right). **E.** *Egr-1* mRNA levels in primary neurons. Results (Means ± SEM; n=3; N=3) are given as fold induction compared to control the mock. Statistical analysis: one-way ANOVA followed by Dunnett’s multiple comparison test. *ns*: non-significant, *p<0.05.

Intracellular localization in differentiated SH-SY5Y cells has been assessed by confocal microscopy with HA antibodies and with DAPI for the nucleus (Figure 3B). HA-C99 presented a diffuse expression. For the cc-del6 construct, the localization appears mainly associated with the plasma membrane, while the cc-del7 presented predominantly intracellular localization (*punctae*), although plasma membrane localization could also be detected.

Cell surface localization of all the cc-del(X) proteins was further addressed by flow cytometry on living cells using an anti-HA antibody (Supplementary Figure 2). All the constructs could be localized to the plasma membrane. The highest median fluorescence was observed for the cc-del6 construct, in line with confocal microscopy. This confirmed a preferential plasma membrane localization of cc-del6 when compared to other cc-del(X) proteins.

We next examined whether these features, e.g dimeric orientation and differential subcellular localization, also exerted an effect on gene expression in the neuronal cell line (Fig 3C). *Egr-1* expression increased for HA-C99 and for cc-del7, while the expression remained at a similar level to mock infected SH-SY5Y cells for the other constructs.

Finally, we tested whether our model was also relevant for rat primary cortical neurons. Primary neurons were transduced with lentiviruses expressing mock vector, HA-C99, cc-del6 and cc-del7. The protein expression profile was examined by immunoblotting with the Cter APP antibody and we observed that cc-del7 produced more AICD in primary neuronal cultures (Figure 3D). We next assessed gene regulation upon cc-del7 expression in primary neuronal cultures. The RNA was extracted and monitored by qPCR to measure *Egr-1* gene expression (Figure 3E). The results show a significant increase in mRNA expression with HA-C99 and cc-del7 compared to the mock, confirming their gene regulation activity.

### cc-del6 favors the production of SDS-resistant oligomers by γ-secretase processing

So far, the cc-del7 construct displayed unique features compared to other cc-del(X) proteins by promoting the release of transcriptionally active AICD and gene expression. Apart from AICD, the processing of APP by γ-secretase produces a class of Aβ peptides directly involved in senile plaque formation. To assess in detail the amyloidogenic profile of all the cc-del(X) fusion proteins, CHO cells transiently expressing cc-del(X) constructs were lysed and separated by SDS-PAGE, then analyzed by Western blotting. The constructs were observed as monomers at the expected size (14-15 kDa) with the Cter antibody. Additional bands were detected at a lower size corresponding likely to a C83-like CTF (Figure 4A). Strikingly, the anti-HA antibody also detected a band around 28 kDa and 60 kDa uniquely for cc-del6. As we did not observe these bands with the Cter APP antibody, we infer the identities of these bands to be oligomeric “Aβ-like” species that we call “oligo*” or “oligo**” corresponding to the 28 kDa and 60 kDa, respectively. Importantly, expression of HA-C99 also led to the appearance of low levels of SDS-resistant 28 kDa oligomers, and those were also only recognized by the anti-HA antibodies.

**Figure 4.**
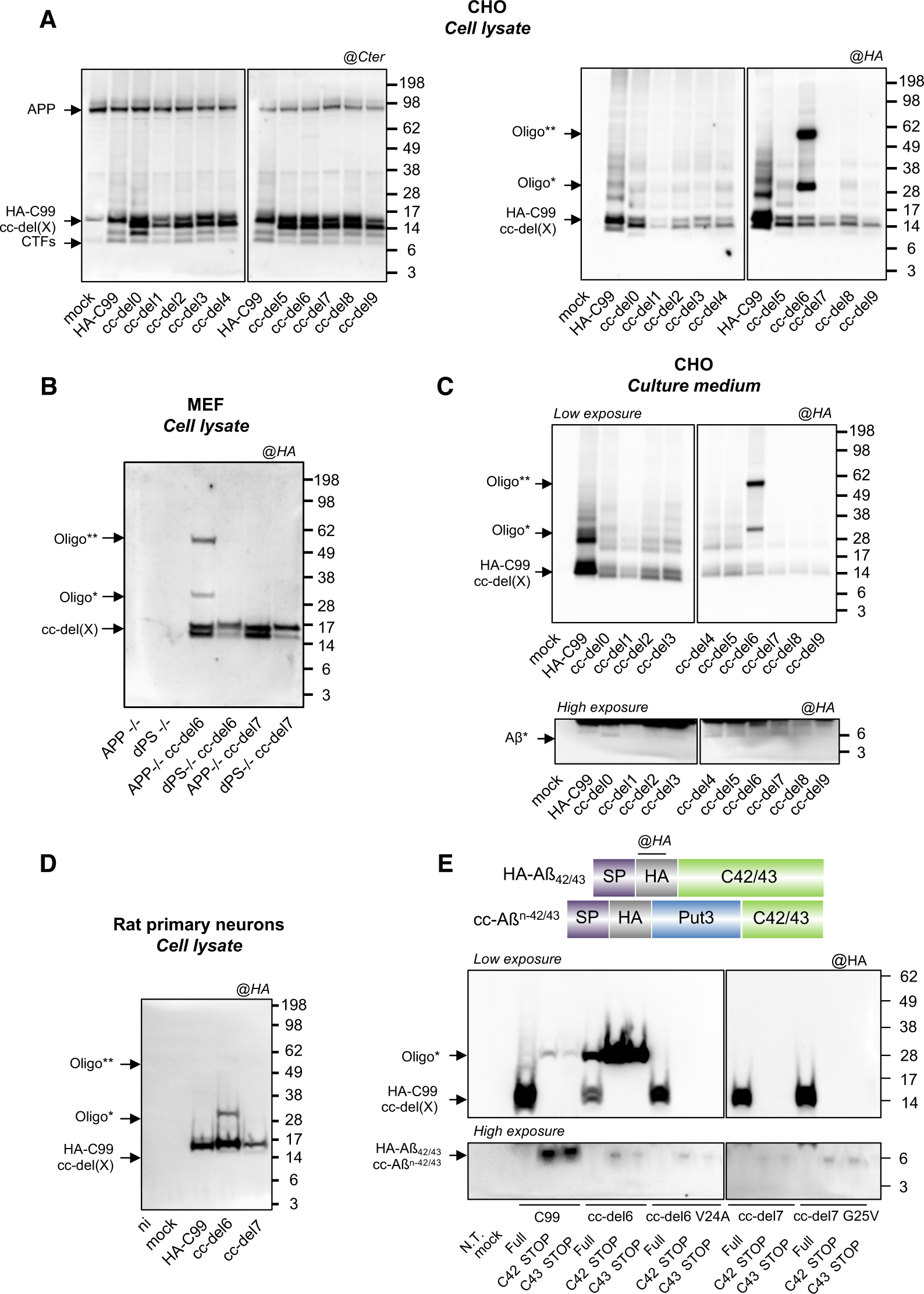
cc-del6 orientation in conjunction with pro-β motif favors the production of SDS-resistant oligomers by **γ**-secretase processing. **A.** Analysis of the profile of cc-del(X) constructs in denaturing conditions. Cells lysates were analyzed by Western Blotting and revealed with APP Cter antibody (left) or HA antibody (right). **B.** Western blot of MEF APP+/+, MEF double knock out for presenilin 1 and 2 (dPS-/-). Detection was performed using the HA-tag antibody. **C.** Analysis of extracellular media. Immunoprecipitation was performed on the supernatant. CHO cells were expressing HA-C99 and cc-del(X) constructs, analyzed by Western blotting and revealed with HA antibody (top). At high exposure (bottom) monomeric Aβ-like (Aβ*) was detected. **D.** Neuronal lysates were analyzed 4 days after infection by Western blotting and revealed with HA antibody. **E.** Top: Representation of HA-C42/43 and cc-A*β*^n-42,43^ constructs. Bottom: Analysis of the profile of truncated forms (C42/C43) of cc-del(X) constructs and Aβ-like peptides (top) in denaturing conditions. Cell lysates were analyzed by Western Blotting and revealed with HA antibody (bottom).

To analyze whether the above-mentioned oligomeric bands were produced by γ-secretase activity, we generated cell lines stably expressing cc-del(X) in murine embryonic fibroblast cells (MEFs) deficient for presenilin 1 and presenilin 2 (MEF dPS-/-), which are devoid of γ-secretase activity. The expression profile of cc-del(X) constructs was identical in MEF APP-/- cells (Figure 4B) and CHO cells (Figure 4A), with only MEF APP-/- cc-del6 exhibiting oligomeric bands around 28 kDa and 60 kDa. We compared the expression profiles of cc-del6 in MEF APP-/- and MEF dPS-/- cells. No oligomeric bands were observed in MEF dPS-/- cc-del6 cells, indicating that processing by the γ-secretase is mandatory for production of the SDS-resistant 28 kDa and 60 kDa oligomeric bands.

We further investigated whether oligomers detected in cell lysates are also released in the culture medium. The release of cc-del(X) fragments was measured in the supernatant of CHO cells by immunoprecipitation with anti-HA antibodies. We observed the presence of the two bands at 28 kDa and 60 kDa secreted in the supernatant for cc-del6 (Figure 4C) and also the presence of cc-del(X) monomers. Importantly, upon higher exposure, we noticed for all the cc-del(X) constructs the presence of a band around 6 kDa. This band likely corresponds to the monomer of the Aβ-like protein (Aβ*) produced by the cc-del(X) constructs.

We next assessed whether these Aβ-like oligomers could also be detected in primary neurons transduced with lentiviruses expressing HA-C99, cc-del6 and cc-del7. Immunoblotting with the HA antibody (Figure 4D) showed that cc-del6 is able to form oligomers in primary neuronal culture, like in the other cell lines tested.

### The production of SDS-resistant oligomers requires a pro-β motif and a specific orientation of Aβ-like peptides

CHO cells were also transfected with the constructs without the Put3 coiled coil system, differing only by deletions in the N-terminus of C99 (Supplementary Figure 3A). This was performed in order to investigate whether the oligomerization profile of C99 is impacted by the deletions. All constructs up to del5 expressed SDS-resistant oligomers in cell lysates (Supplementary Figure 3B). Further deletions (del6-9) impaired insertion in the membrane as the TM segment became too short (unlike the Put3 fusions which always contained sufficient Leu residues) and did not give rise to any oligomers. These results indicate that progressive deletions at the outset of the TM domain do not by themselves induce oligomerization and that Put3 by providing the required hydrophobicity allows membrane insertion, cell-surface localization, in addition to imposing precise orientations.

The expression of cc-del(X) in dimeric form was verified by transient transfection in CHO cells by blue native PAGE with Cter APP antibody and the presence of oligomers with HA antibody (Supplementary Figure 3C). The presence of bands at the expected size around 20 kDa (C-ter antibody) confirmed that all cc-del(X) fusion proteins are able to form dimers. We also observed (with the HA antibody) the presence of bands around 66 kDa for most constructs with highest levels in cc-del3, cc-del5 and cc-del6. Those results, correlated to the previous results (Figure 4A), show that although all the constructs are detected as dimers in native conditions, only cc-del6 is able to form SDS-resistant oligomers, oligomers induced by other constructs, and notably cc-del3 and cc-del5, are not SDS-resistant.

Next, we used the Tango algorithm to assess whether the addition of Put3 could provide an environment favoring the production of the SDS-resistant oligomers observed upon cc-del6 processing. The Tango algorithm predicts the cross-β aggregation propensity of amino acids sequences (referred to as pro-β motif) based on physico-chemical principles. Its accuracy was previously validated on a set of 250 peptides including the amyloid beta peptide (Fernandez-Escamilla et al., 2004). As expected, the C-terminal segment of the different forms of Aβ-like peptides was predicted to have a high propensity for cross-β aggregation in all cc-del(X) constructs (Supplementary Figure 4A). In cc-del0 to cc-del2, all or part of the ^16^LVFF^20^ motif (due to the single amino acid deletion) was also predicted to have a high propensity for cross-β aggregation (Supplementary Figure 4A), in agreement with previous studies (Fernandez-Escamilla et al., 2004). Strikingly, cc-del6 was the only construct beyond cc-del2 to contain a pro-β motif, ^20^ALLLV^24^, according to the algorithm.

The difference between cc-del6 and cc-del7 was the Val at position 24 present in cc-del6 but deleted in cc-del7. We mutated Val24 in cc-del6 to Ala and Gly25 to Val in cc-del7 (Supplementary Figure 4B) to assess whether this motif (ALLLV) was key for the production of SDS-resistant oligomers. The mutations were predicted to reduce cross-β aggregation propensity in cc-del6 and increase cross-β aggregation in cc-del7 without changing their dimeric TM orientation (Supplementary Figure 4C). Strikingly, the V24A mutation in cc-del6 led to the total disappearance of SDS-resistant oligomers (Supplementary Figure 4D). In contrast, the presence of this pro-β motif in cc-del7 did not result in the release of SDS-resistant oligomers (Supplementary Figure 4D). This motif was thus required but not sufficient for the production of SDS-resistant oligomers.

To discriminate the contribution of different TM orientations of Aβ-like peptides from that of different lines of cleavage possibly followed by cc-del6 and cc-del7, we created truncated forms of HA-C99, cc-del6 and cc-del7 constructs (with and without the precited mutations) by adding a stop codon at position 42 and 43 (amyloid beta numbering) (Figure 4E, top). As expected, direct transfection of WT C42 and C43 led to the production of SDS-resistant oligomers. The direct transfection of truncated forms of cc-del6 at position 42 and 43 both led to the production of a much larger amount of SDS-resistant oligomers (Figure 4E). In line with our previous results, the V24A mutation blocked the formation of SDS-resistant oligomers in the cc-del6 context (Figure 4E). Importantly, direct transfection of the truncated forms of cc-del7 with or without the G25V mutation did not result in the apparition of SDS-resistant oligomeric bands (Figure 4E). The cc-del6 orientation thus appeared required for the stability of these oligomers.

### cc-del6 and cc-del7 orientations favors distinct processing lines by tripeptide cleavage by γ-secretase

A key goal was to assess whether different dimeric TM orientations of C99 favored distinct processing lines. We first created a specific protocol to identify the composition of the oligomers produced upon cc-del6 processing by mass spectrometry analysis (Supplementary Figure 5A). The oligomeric bands observed from the medium after IP (Supplementary Figure 5A arrow 1) can be disrupted with formic acid (Supplementary Figure 5A arrow 2). The eluates obtained after IP with formic acid were analyzed by mass spectrometry (Supplementary Figure 5A arrow 3). The oligomeric bands were composed of Aβ* that contained residues 40 and 43 of the Aβ sequence at the C-terminus (C40 and C43, respectively) (Figure 5A). Mass spectrometry data are shown in Supplementary Figure 6. Next, we assessed whether our fusion proteins obeyed the tripeptide cleavage model of γ-secretase and that cc-del6 thus followed the processing line Aβ_49_>Aβ_46_>Aβ_43_>Aβ_40_. According to Bolduc et al. (2016), presenilins contain three putative S’ pockets that can each accommodate one amino acid at a time and occupancy of these three pockets precedes cleavage of a tripeptide. The S1’ and S3’ pockets are large and can easily accommodate any amino acids. The smaller S2’ pocket is less prone to accommodate large aromatic residues like phenylalanine. In consequence, placing an aromatic residue at a given position is predicted to reduce overall cleavage beyond this point, although it was shown that γ-secretase was able to skip multiple residues to cleave at a further position (Bolduc et al., 2016). If cc-del6 follows the tripeptide cleavage model, mutating the amino acids at position 44 and 45 to phenylalanines would decrease further processing and result in accumulation of C46. Similarly, preventing initial ε-cleavage by mutating the residues at position 50-51 to phenylalanine should reduce overall processing.

**Figure 5.**
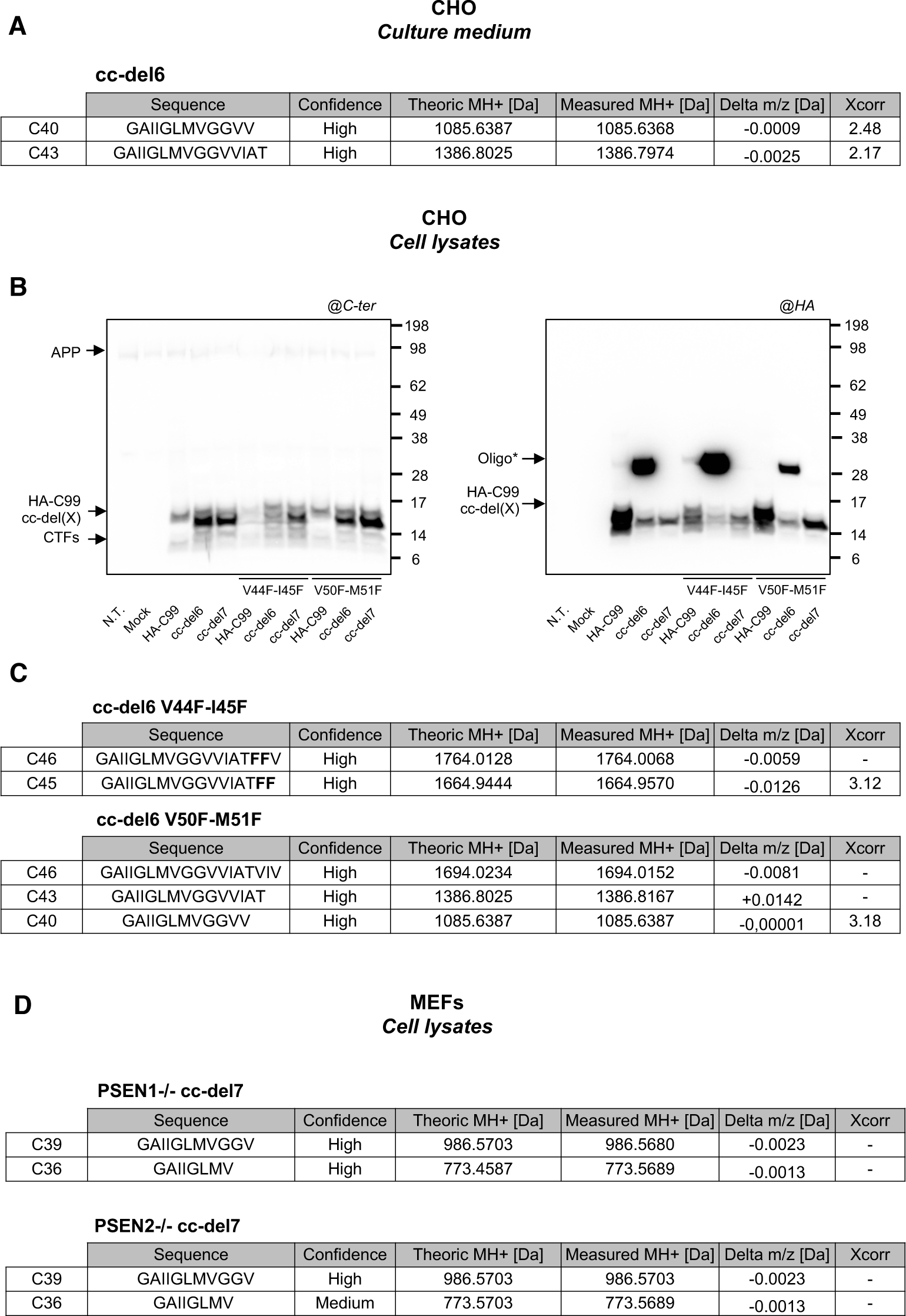
Processing lines favored by cc-del6 and cc-del7 orientations. **A.** Mass spectrometry analysis of oligomers from culture medium produced upon cc-del6 processing after disruption with formic acid. The results are summarized in the table. XCorr values are given for MS/MS spectra identified by Sequest. Spectra are available in Supplementary Figure 6. **B.** Analysis of the profile of HA-C99, cc-del6 and cc-del7 constructs with or without the V44F-I45F and V50F-M51F mutations in denaturing conditions. Cells lysates were analyzed by Western Blotting and revealed with APP Cter antibody (left) or HA antibody (right). **C.** Mass spectrometry analysis of oligomers from cell lysates produced upon cc-del6 processing with the V44F-I45F or V50F-M51F mutations after disruption with formic acid. The results are summarized in the table. XCorr values are given for MS/MS spectra identified by Sequest .Spectra or PRM traces are available in Supplementary Figure 7 and 8. **D.** Mass spectrometry analysis of Commassie Blue positive bands detected in PSEN1 -/- or PSEN2 -/- MEFs infected with cc-del7. The results are summarized in the table. PRM traces results are available in Supplementary Figure 9.

In agreement with these predictions, we observed decreased production of SDS-resistant oligomers in presence of the V50F-M51F mutation in cc-del6 and absence of such oligomers in HA-C99 (Figure 5B). In contrast, the V44F-I45F mutation led to increased accumulation of SDS-resistant oligomers in HA-C99 and even more importantly in cc-del6 (Figure 5B). In order to identify the composition of these oligomers produced upon cc-del6 processing, we used the protocol described above for mass spectrometry analysis of cell lysates after IP (Supplementary Figure 5B). Optimization of the detection enabled the identification of the C-terminus of C46, C43 and C40 in the lysates of cells expressing cc-del6 V50F-M51F (Figure 5C). In contrast, the band observed in cells expressing cc-del6 V44F-I45F was mainly composed of C46 containing the V44F-I45F mutation. We also detected a small amount of C45 with the V44F-I45F mutation (Figure 5C and Supplementary Figure 7), which can be interpreted as a side effect of the substitution to phenylalanines. Mass spectrometry data are given in Supplementary Figure 7 and 8. Together, these results give strong evidence that cc-del6 obeys the tripeptide cleavage of γ-secretase and that it follows the Aβ_49_>Aβ_46_>Aβ_43_>Aβ_40_ processing line.

We also investigated the processing of cc-del7. We reasoned that if the cc-del6 orientation promoted the line of cleavage starting at position 49, a rotation of 110° counter-clockwise, like in cc-del7, bringing the more distal ^33^Gly-x-x-x-Gly^37^ in the interface, would lead to less tilt and thus an initial cleavage upstream at position 48 and thereby to the Aβ42 processing line. cc-del7 produced oligomers based on native gels, but the instability of Aβ-like peptide oligomers in SDS did not allow the acquisition of conclusive results by mass spectrometry analysis with a similar protocol as the one used for cc-del6, although suggestive data indicated the possibility of the Aβ48 line of cleavage.

To enhance the chance of purifying cc-del7–derived Aβ-like peptides and possibly aggregates/oligomers of cc-del7 products, we used MEFs that are knock-out for either PSEN1 or PSEN2 that we infected with cc-del7. It has been shown that in such MEFs, compensatory overexpression of the remaining PSEN occurs (Stanga et al., 2018), possibly triggering the processing of APP substrates. In addition, a recent study showed that longer Aβ forms (Aß42) were preferentially produced by PSEN2 in endocytic compartments to generate an intracellular Aβ pool (Sannerud et al., 2016). This observation is fitting with our data indicating that Aβ-like peptides produced by cc-del(X) processing accumulate in cell lysates. In addition, PSEN1 and PSEN2 have been involved in autophagy and lysosomal proteolysis (Lee et al., 2010; Neely et al., 2011); their absence thus favors the accumulation of cleavage products that would normally be degraded by these pathways. Confocal microscopy also showed large accumulation of HA positive intracellular vesicles in SH-SY5Y cells infected with cc-del7 (Figure 3B), suggesting that cc-del7 associated oligomers are present in stress granules that tend to accumulates when phagocytosis is impaired.

We detected large accumulation of aggregated proteins by Ponceau red and by Coomasie Blue staining in MEFs PSEN1 or PSEN2 KO infected with cc-del7. These oligomeric bands were relatively stable in NP40, but sensitive to SDS unlike cc-del6 oligomers. We analyzed the Coomasie blue positive band, ranging from 49-62 kDa and recognized by HA antibodies, by mass spectrometry. We detected with high levels of confidence C36 and C39 peptides in these bands. More specifically, mass spectrometry showed the presence of C39 and C36 in PSEN1 -/- cc-del7 and C39 in PSEN2 -/- cc-del7. These fragments were not detected in a band of the corresponding size from non-infected MEFs. In line with our hypothesis, mass spectrometry analysis also detected the presence of p62, a marker of autophagosome, in Coomassie Blue positive bands in both PSEN1 and PSEN2 KO MEFs infected with cc-del7 but not in negative control. Together, these results indicated that products from the Aβ_42_>Aβ_39_>Aβ_36_ processing line are detectable in cells expressing the cc-del7 construct.

### A model for the choice of γ-secretase cleavage lines as a function of dimeric orientation

A model of the dimeric orientations adopted by cc-del7 and cc-del6 based on the solved structure of the TM domain of APP (pdb: 2LOH) and modified according to (Sato et al, 2009; Zhou et al, 2019) is shown in Figure 6. We hypothesize that dimerization via the Gly-x-x-x-Gly motifs will affect the accessibility and configuration for initial binding of γ-secretase on dimeric C99 by exposing ε-sites and, subsequently, γ-sites for AICD and Aβ release, respectively. Specifically, as a function of which Gly-x-x-x-Gly motif is in the interface, the tilt of the helices will differ. Crossing at the N-terminus of the TM domains (for cc-del6) with Gly25 and Gly29 in interfacial “d” and “a” positions will result in a higher tilt at the C-terminus, when compared to crossing in the middle of the TM domain (for cc-del7). A higher tilt will lead to a lower position exposed, at the ε-cleavage site. After initial binding, the dimeric substrate would be uncoupled to allow processing.

**Figure 6.**
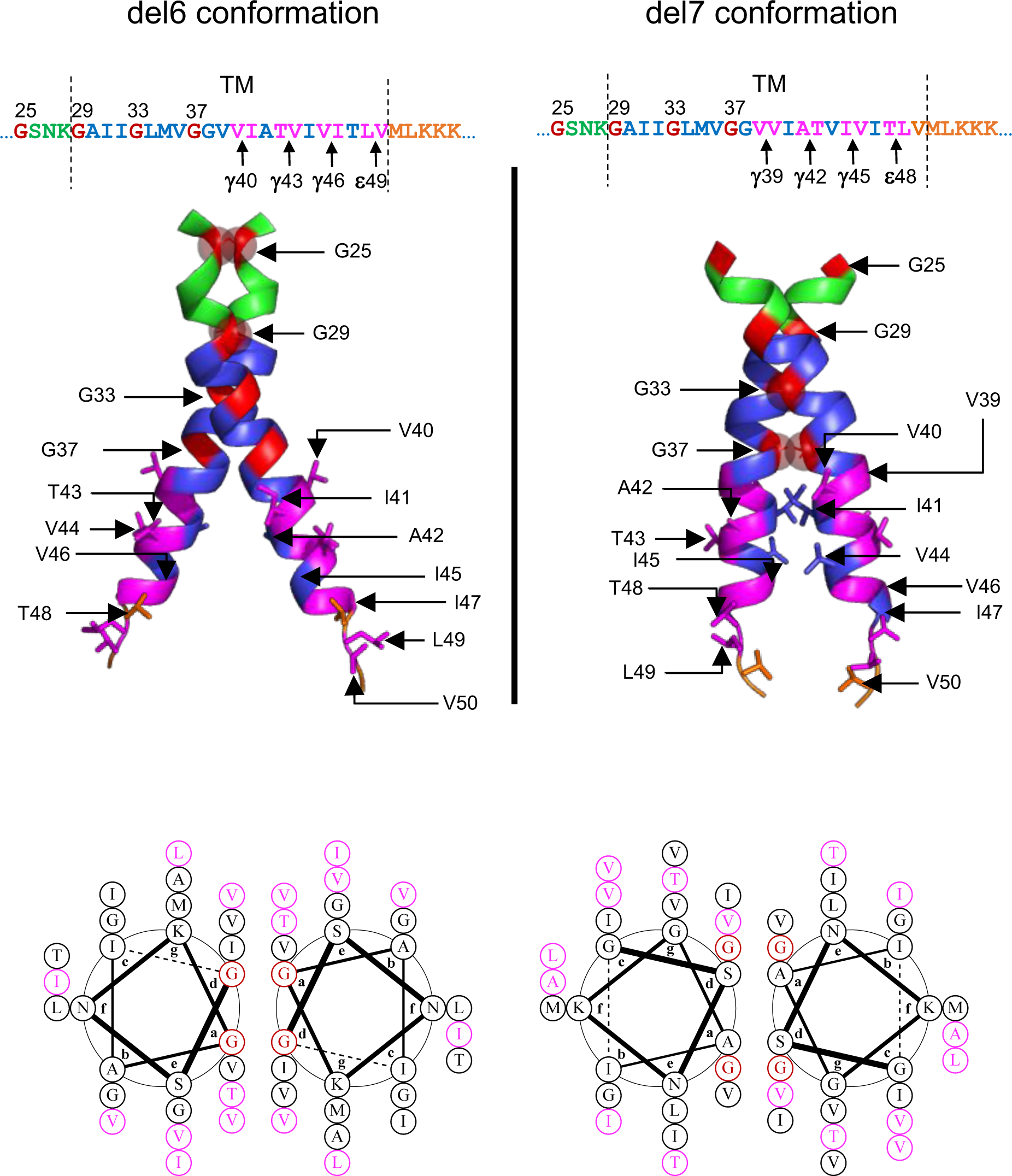
Dimeric helical structure cc-del6 and C99 cc-del7 Top: Dimeric model structures of cc-del7 and cc-del6 fusion proteins were adapted base on pdb: 2loh (Nadezhdin et al, 2012). The juxtamembane (JM) region is in green, transmembrane region (TM) in blue and AICD in orange. Glycine residues from GxxxG motifs are in red and the different positions of γ-secretase cleavage in pink. The arrows indicate the amino acids in the helix (Aβ numbering). Bottom: Helical wheel diagrams of the cc-del7 and cc-del6 fusion proteins. Glycine residues in the interface are in red and the residues accessible for γ-cleavage are in pink. The model does not take into account potential distortions/kinks induced by ^33^GlyxxxGly^37^Gly^38^.

In Figure 6 (left panel), with the helical lateral view, we show that the C99 cc-del6 dimer contains the ^25^Gly-x-x-x-Gly^29^ motif in the interface, which forms a more upstream crossing interface that requires a higher tilt of the helices. This exposes the amide bond of an amino acid further down the helix, at or after residue 49 within the ε-site. At those positions, the helix is already unraveled, as we have proven for our constructs as well with the L49C cross-linking (Supplementary Figure 1). Cleavage at residue 49 would release AICD and process upstream at the γ43 and γ40 sites for releasing Aβ43 and Aβ40-like peptides respectively (Qi-Takahara et al, 2005; Takami et al, 2009). The model that we propose fits very well with the experimental data in cell free assays for AICD production and mass spectrometry for the Aβ isoforms production, where we see for cc-del6 a preferential cleavage at C46, C43 and C40.

In Figure 6 (right panel), with the helical lateral view, we show that C99 cc-del7 contains ^33^Gly-x-x-x-Gly^37^ in the interface with Gly33 and Gly37 in interfacial “d” and “a” positions. Our data in MEFs deficient for PSEN1 or PSEN2 suggest that initial cleavage could occur at residue 48. This residue would be preferred in this more parallel dimer. Following tripeptide cleavage by γ-secretase, such initial cleavage at ε-site 48 would favor the Aβ42 processing line and an interesting overlap between signaling and generation of Aβ42 could be invoked.

## Discussion

Our main observations are that C99 TM domain dimerization affects its cleavage by γ-secretase, and that different functional outcomes in terms of signaling and Aβ oligomerization depend on two of the seven possible symmetric dimeric orientations of the C99 TM domain, before and after processing. We have employed coiled coils to impose precise dimeric interfaces. We validated the predicted interfaces by Cys-cross-linking. We found that one dimeric interface (cc-del7) leads to enhanced release of transcriptionally active AICD and *Egr-1* gene expression as well as its downstream target *Bace1*. This orientation favors initial cleavage at position 48 and thereby the Aβ42 processing line. Another orientation, cc-del6 generates Aβ-like peptides that form SDS-resistant oligomers. Cleavage of cc-del6 at residue 49 leads to formation of SDS-resistant oligomers ending at positions 43 and 40 of Aβ. SDS-resistance of these oligomers requires the presence of a pro-β motif in addition to a precise dimeric orientation of Aβ-like peptides.

The two interfaces are related and rotated relative to each other by 110°. cc-del7 forms a dimer that crosses the TM helices at the middle ^33^Gly-x-x-x-Gly^37^ TM motif, while cc-del6 crosses the helices upstream at the ^25^Gly-x-x-x-Gly^29^. While both are processed by a presenilin-dependent γ-secretase activity, our data and model argue that the initial cleavage at the ε-site differs because the tilt of helices differs exposing different sites to the γ-secretase, the higher the tilt in cc-del6 dimer, the lower start of cleavage (at 49).

Importantly, the cc-del7 fusion protein exhibits the same dimeric interface as cc-del0, but with a shorter extracellular region. It has been reported that the ^16^LVFF^20^ motif forms a β-sheet that prevents processing of C99 by γ-secretase (Hu et al, 2017; Tian et al, 2010). This particular motif is absent in the cc-del7, but present in cc-del0. Indeed, no effect has been observed in term of signaling for the cc-del0, consistent with decreased processing. Together, it indicates that the conformation of cc-del7 TM dimers is the active one for APP nuclear signaling.

The precise regulation of APP amyloidogenic processing depends on its subcellular localization and the nature of the γ-secretase present in the cellular compartments (presenilin 1- or presenilin 2-dependent). Our confocal immunofluorescence data from human neuroblastoma cell lines (Figure 3B) indicated that cc-del7 appears localized to intracytoplasmic structures, while cc-del6 was predominantly localized to the plasma membrane. The processing of cc-del6 and cc-del7 may occur in different cellular compartments, with implications on the fate of produced Aβ or AICD. Recently, it has been reported that amyloidogenic processing by presenillin 2 occurs preferentially in recycling endosomes and late endosomes/lysosomes compartments (Sannerud et al, 2016), which would fit to the localization of our cc-del7 dimer.

The comprehensive model that we propose is C99 dimers with an orientation where ^33^Gly-x-x-x-Gly^37^ is in the interface (cc-del7) allow high levels of trimming by γ-secretase likely starting at position 48 (Figure 6). A mechanism has recently been described where stable substrate binding to γ-secretase is important for correct processing (Szaruga et al, 2017). This binding initially occurs just upstream of the region we described to shift from the TM helical structure to random coil (Sato et al, 2009). After initial binding, γ-secretase would induce unravelling of the helix for initial cleavage at the ε-site before cutting upstream in sections of three amino acids, to form gradually shorter Aβ species (Bolduc et al, 2016). Our mass spectrometry data in MEFs PSEN1-/- and PSEN2 -/- suggest that initial cleavage occurs at residue 48 with tripeptide cleavage then releasing Aβ45, Aβ42, Aβ39 and, in some cases, Aß36. The fact that Aβ-like peptides ending at position 42 were hardly detectable in our set up probably indicates that the carboxypeptidase (trimming) activity of the γ-secretase was particularly efficient to produce shorter peptides in the cc-del7 conformation..

In pathological conditions, it is possible that the generated Aβ peptides, like Aβ42, might then change orientation by rotating clockwise 110° to the same orientation where ^25^Glyx-x-x-Gly^29^ motif would come into the interface, corresponding to the orientation of cc-del6, and form stable Aβ oligomers. The shift from cc-del7 to cc-del6 would result in shifting the helix crossing point from G^33^xxxG^37^ closer up to the extracellular juxtamembrane region at ^25^Glyx-x-x-Gly^29^. Such an upward shift might alter interactions with membrane lipids (Zhou et al, 2019) and favor SDS resistance of the released Aβ peptides, just like in the case where the initial cleavage would occur at or after residue 49. The proportions of C99 that adopt one or the other dimeric interfaces might depend on their precise intracellular localization, membrane composition, pH, and temperature, as the precise interaction between the γ-secretase and its substrates, especially intermediates between Aβ49 and Aβ40 were shown to depend on temperature (Szaruga et al, 2017).

While our study utilizes constructs with TM-cytosolic domain identical to those of C99, the N-terminus of our constructs codes for the Put3 coiled coil. This is a limitation of our study. The Aβ-like peptides derived from cc-del6 processing are specifically able to generate SDS resistant oligomers of ∼28 kDa and ∼60 kDa (Figure 4) referred to as oligo* and oligo**, respectively, but these are Aβ-like peptides as they contain a segment of the Put3 coiled coil at their N-terminus. Because oligomers are nucleated by the coiled coil helices, we suggest the possibility that such oligomers might adopt transient α-helical structure during initial nucleation. We showed that SDS resistance is reminiscent of the similar resistance of the glycophorin A TM dimer (Bormann et al, 1989), and requires: i) close apposition of the mediated by close apposition of a specific ^25^Gly-x-x-x-Gly^29^ motif in the interface and ii) a pro-β motif for secondary structure stabilization as suggested by the work of Misra et al. (2016). The fusion of Put3 coiled coil with the TM-cytosolic domain of C99 generates a novel sequence where Val24 of C99 is key for oligomerization, allowing the reconstitution of a pro-β motif. We posit that the role of this motif is reminiscent of this of the ^16^LVFF^20^ motif that we and others previously showed to be key for the processing and adopting a specific secondary structure that prevents amyloidogenic processing (Hu et al, 2017; Tian et al, 2010). Importantly, we show that substitution of Val24 in the pro-β ^20^ALLLV^24^ motif is sufficient to impede SDS resistance of Aβ-like peptides ending at position 42 and 43 in the cc-del6 context. This is line with previous findings that substitution of Val17 (among others) in the ^16^LVFF^20^ motif similarly impedes SDS-resistance (Hilbich et al., 1992). Interestingly, SDS-resistant Aβ oligomers were detected in brains from Alzheimer’s patients (Roher et al., 1996).

According to our model and to the cross-linking data in Supplementary Figure 1, the cc-del6 construct exhibits a particular orientation in which two upstream glycines, Gly25 and 29 of the ^25^Gly-x-x-x-Gly^29^ motif are in the interface in “d” and “a” positions. Thus, we suggest that while these Gly residues mediate helix crossing before cleavage, they might also mediate initial nucleation of oligomers after cleavage. Still, we cannot nominally ascribe to them an *α*-helical structure without further studies, as they might switch to other secondary structures, as suggested by the data described in Figure 4E and the work of Misra et al. (2016). The observation that the other cc-del(X) fusion proteins exhibiting different orientations also dimerize (Supplementary Figure 3C), produce Aβ-like monomers, but do not lead to the formation of SDS-resistant oligomers shows that the precise dimeric TM orientation with a particular Gly-x-x-x-Gly motif in the interface is key for the production of stable Aβ-like oligomers.

We show that cc-del6 obeys the tripeptide cleavage by γ-secretase, with processing likely starting at position 49, trimming in sections of three residues occurs (Bolduc et al, 2016), allowing the formation of Aβ46, −43 and −40. Several studies have shown that APP amyloidogenic processing leads to higher Aβ40 production compared to Aβ42 (10:1 ratio), but one study has demonstrated that Aβ43 is more common than Aβ40 in amyloid plaque cores found in AD brains (Welander et al, 2009). The aggregation propensity and potential pathogenic role of Aβ43 have been suggested (Saito et al, 2011; Veugelen et al, 2016) and recently documented (Szaruga et al, 2017). We suggest that different TM orientations can result in different lines of cleavage but that production of stable, SDS-resistant oligomers is linked with the specific TM orientation imposed in cc-del6 together with the presence of a central pro-β motif such as the ^20^ALLLV^24^ present in cc-del6 sequence or the ^16^LVFF^20^ motif of the human Aβ sequence.

The role of the precise dimerization of the C99 TM dimerization in signaling and processing might open novel therapeutic avenues to limit amyloidogenic processing. These results might lead to the identification of the physiologic targets of APP signaling, which remain poorly known as does the physiological ligand for APP and related proteins APLP1 and APLP2. One provocative scenario is that in aged individuals a higher degree of signaling by APP is required to compensate for age-related hypoxia and senescence. Such overuse of the pathway modelized by cc-del7 would drive formation of pathogenic amyloidogenic peptides and possibly induce a self-perpetuating amyloidogenic loop by increasing the expression of *Egr1* and its downstream target *Bace1*. We suggest that our fusion protein cc-del7 could become a model of constitutive active APP or C99 useful for identification of physiologic APP signaling targets. Finally, this model would also explain pathologic signaling by Familial cases of Alzheimer’s disease (FAD), where certain mutations favor dimerization via the Gly-x-x-x-Gly motifs like cc-del7 (Tang et al, 2014). Such mutations were recently also reported as somatic acquired mutations in sporadic Alzheimer’s cases, thus increasing the relevance of understanding the mechanisms behind the pathogenicity of FAD mutations (Lee et al, 2018).

## Materials and Methods

### cDNA constructs

Wild type C99 with a HA-tag was cloned into the pLVX-CMV bicistronic retroviral vector. cDNAs coding for the ten fusion proteins cc-C99^18–99^ to cc-C99^27–99^ (referred to cc-del0 to cc-del9) and their associated mutants were generated by overlap extension using PCR with synthetic nucleotides. All constructs were sequenced by Macrogen.

### Primary neuronal cultures

Primary cultures of cortical neurons were prepared from E18 rats embryos. The cortex was dissected and dissociated in HBSS depleted of calcium and magnesium. HBSS (with calcium and magnesium) was added and the mixture centrifuged through Fetal Bovine Serum (FBS) for 10 min at 1000g to pellet cells. Cells were plated at 1.6 x 10^6^ cells/cm² in culture dishes pre-treated with 10 µg/ml of poly-L-lysine in phosphate buffered saline (PBS) and cultured for 7 days *in vitro* in Neurobasal® medium supplemented with 2% v/v B-27® supplement medium and 1mM L-glutamine (all from Life Technologies) at 37 °C, 5% CO_2_ in a humidified incubator. Half of the medium was renewed every 2-3 days prior to infection.

### Cell cultures and transfection

Chinese hamster ovary (CHO) cell lines (ATCC) were grown in Ham’s F12 medium (Thermofisher Scientific) and were supplemented with 10% of FBS (Sigma-Aldrich). The MEF (Murine Embryonic Fibroblast) APP-/- cell lines, MEF dPS-/- (presenilin 1 and 2 knock out) and the human neuroblastoma SH-SY5Y cell lines (ATCC) were grown in DMEM/F12 medium (Thermofisher Scientific) and were supplemented with 10% of FBS. SH-SY5Y cells were differentiated into neurons with 10 μM of retinoic acid (Sigma-Aldrich) after 7 days. All cell cultures were maintained at 37 °C in a humidified incubator (5% CO_2_). For transient transfection, cells were seeded at a density of 3.10^5^ cells/well 24h before transfection (2 μg of pLVX-CMV DNA/well) with lipofectamine 2000™ according to the manufacturer’s instruction (Invitrogen). The SH-SY5Y were treated with 300 nM of trichostatin A (TSA) during 24h after differentiation prior to RNA extraction.

### Cross-linking studies

The ten cc-del(X) fusion proteins were mutated by directed mutagenesis (Gly29 into Cys) to generate the cc-del(X) G29C constructs. After transfection, CHO cells were washed in PBS and resuspended in 1 ml of PBS containing 1 mM MgCl_2_ and 0.1 mM CaCl_2_. Cross-linking was performed with a final concentration of 100 μM of the crosslinker N, N’-1, 2-phenylenedimaleimide (o-PDM) dissolved in DMSO for 10 min at room temperature. Cells were lysed in NP40 lysis buffer containing 2% of β-mercaptoethanol to quench the reaction. Lysates were analyzed by Western blotting, with an antibody directed against the C-terminus fragment of APP (Cter).

### Generation of cell lines

MEF and SH-SY5Y cell lines stably expressing the different cc-del(X) fusion proteins were established by viral transduction. Viruses were generated by transient transfection in HEK cells with the vectors psPAX2 and pMD2.G (Clontech) using the calcium phosphate method, according to the manufacturer’s instruction (Promega). Viruses were then concentrated with Centricon filter units (Merck Millipore). Infected cells were selected with 20 μM of puromycin for 1 week and maintained in culture with 2 μM of the antibiotic.

### Western blotting and blue native gel electrophoresis

For SDS/LDS-PAGE electrophoresis, cells were commonly lysed in Laemmli 2X buffer. For isolation of Coomassie Blue positive bands associated with cc-del7, Laemmli lysis buffer was replaced with NP-40 lysis buffer. LDS sample buffer (ThermoFisher Scientific) was added to the lysates and the mix was boiled 10 minutes at 70°C. Proteins (25 µg) were separated on Bolt™ 4-16% Bis-Tris Protein Gels (Invitrogen) and transferred onto nitrocellulose membrane via iBlot gel transfer stacks (Invitrogen). For Blue Native PAGE, cells were lysed in native blue buffer, DDM 10% (Invitrogen) and 100X protease inhibitor (Pierce). After centrifugation (100,000*g*, 1 h at 4 °C) supernatants were collected (15 µg of proteins) and 5% of sample buffer added. Protein lysates were separated using the Novex® NativePAGE™ Bis-Tris gel system (Invitrogen). The electrophoresis and transfer were performed as per manufacturer’s instructions. Membranes were incubated overnight at 4 °C with primary antibody at following dilutions: 1:1,000 anti-HA, [3F10] (Roche); 1:2,500 anti-Amyloid Precursor Protein [Y188] (referred to Cter - Abcam); 1:5,000 anti-Amyloid β, WO-2 MABN10 (Merck Millipore). Membranes were washed, and further incubated with the secondary antibody (1:5,000 anti-rat [7077S]; anti-rabbit [7074S]; anti-mouse [7076S] (Cell Signaling Technology)) conjugated to horseradish peroxidase followed by ECL (Thermo Scientific). Detection of membranes was performed with the Vilber Fusion Solo S system.

### Immunoprecipitation and immunoblotting

CHO cells were transfected with HA-C99 and cc-del(X) expression plasmids. The supernatant was collected, lyophilized, resuspended in water and pre-cleared with recombinant protein G agarose (Invitrogen). Immunoprecipitation of lysates was performed after lysis in NP-40 lysis buffer. Immunoprecipitation was performed with a monoclonal anti-HA antibody (Roche). Immunoblotting was performed as described above, using a different HA-Tag antibody (C29F4 Rabbit mAb #3724 - Cell Signaling Technology) as primary antibody.

### Mass spectrometry

For the identification of peptides, samples were reduced and alkylated before digestion O/N with trypsin at 30 °C in 50 mM NH_4_HCO_3_ [pH 8.0]. Peptides were dissolved in solvent A (0.1% TFA in 2% ACN), directly loaded onto reversed-phase pre-column (Acclaim PepMap 100, Thermo Scientific) and eluted in backflush mode. Peptide separation was performed using a reversed-phase analytical column (Acclaim PepMap RSLC, 0.075 x 250 mm, Thermo Scientific) with a linear gradient of 4%-36% solvent B (0.1% FA in 98% ACN) for 36 min, 40%-99% solvent B for 10 min and holding at 99% for the last 5 min at a constant flow rate of 300 nl/min on an EASY-nLC 1000 UPLC system. The peptides were analyzed by an Orbitrap Fusion Lumos tribrid mass spectrometer (ThermoFisher Scientific). The peptides were subjected to NSI source followed by tandem mass spectrometry (MS/MS) in Fusion Lumos coupled online to the UPLC. Intact peptides were detected in the Orbitrap at a resolution of 120,000. Data-dependent ions were selected for MS/MS using HCD setting at 35; ion fragments were detected in the Iontrap. The resulting MS/MS data was processed using Sequest HT search engine within Proteome Discoverer 2.3 against a homemade protein database containing the sequences of HA-C99 and cc-del(X) expression plasmids. Trypsin was specified as cleavage enzyme allowing up to 2 missed cleavages, 2 modifications per peptide and up to 2 charges. Mass error was set to 10 ppm for precursor ions and 0.1 Da for fragment ions. Oxidation on Met was considered as variable modification. Peptide spectral matches (PSM) were filtered using charge-state versus cross-correlation scores (Xcorr) and manually validated. For Parallel Reaction Monitoring (PRM) experiments, a targeted mass list was included for the potential C-term sequences, considering potential oxidation of Met and charge state +2. MS1 spectra were obtained at a resolution of 120.000 with an AGC target of 1E6 ions and a maximum injection time of 50 ms, Targeted-MS2 spectra were acquired with an AGC target of 5E4 ions and a maximum injection time of 100ms at a resolution of 30.000.Targeted-MS2 scans were analyzed by Skyline 19.1.0.193 and manually validated.

### Dual luciferase assays

CHO cells were transfected as described above with the following plasmids: cc-del(X) vectors, Fe65, Gal4Tip60 and Gal4RE-luc as reporter gene, according to the procedure previously described (Huysseune et al., 2007). The firefly luciferase activity was standardized with the Renilla luciferase reporter pRLTK. Both luciferase activities were measured with the Dual Glo™ luciferase assay system (Promega).

### AICD detection

MEF APP-/- cc-del(X) cells were grown in 10 cm dishes. Prior to lysis, cells were washed in PBS, collected in cold hypotonic buffer (1 mM MOPS, 1 mM KCl, pH 7.0) and incubated at 25 °C for 25 min. Mechanical lysis was performed by syringe and lysate was clarified at 1,000*g* for 15 min at 4 °C. The post nuclear fraction was centrifuged at 16,000*g* for 40 minutes. Pellets were solubilized in trisodium citrate (150 mM, pH 6.4). Samples were incubated either on ice (considered as time point 0 h) or at 37 °C for 2 h, then analyzed by Western blotting revealed with the APP Cter antibody. Quantification of AICD was performed with Bio-1D software.

### Assessment of **γ**-secretase ability to cleave stable dimers

Cell lines were seeded at a density of 2 x 10^6^ per 10 cm dish for 48 hours. They were washed twice with PBS and re-suspended in PBS with 1mM MgCl_2_ and 0.1 mM CaCl_2_ with 100 µM of the crosslinker N,N’1,2-phenylenedimaleimide (o-PDM) or DMSO (negative control) for 10 minutes at room temperature. Cells were then washed twice with PBS and collected in cold hypotonic buffer (1 mM MOPS, 1 mM KCl, pH 7.0) with 2% β-mercaptoethanol to quench the reaction and 100X protease inhibitor cocktail (Pierce). Lysates were subsequently processed for AICD detection as described above.

### RNA extraction and quantitative real time PCR

Total RNA was purified using RNeasy Mini Kit (Qiagen). Reverse transcription (RT) was performed with the reverse transcriptase core kit using a T3thermocycler (Westburg). Quantitative (q) real time PCR (q-RT PCR) was performed with the MESA Green qPCR MasterMix Plus for SYBR Assay® (Eurogentec) using the Applied Biosystems™ StepOnePlus™ Real-Time PCR System. The relative amplification of cDNA fragments was calculated by the 2^-ΔΔCt^ method with GAPDH as control.

### Surface expression of HA-Tagged cc-del(X) fusion protein

Surface expression of HA-C99 and cc-del(X) fusion proteins was measured with monoclonal HA-antibody (Roche 1:100) by flow cytometry as described previously (Staerk et al., 2011).

### Confocal microscopy

SH-SY5Y cells expressing cc-del6, cc-del7 and HA-C99 were grown on chamber slides with coverglass (ThermoFischer Scientific) and were differentiated over 7 days with retinoic acid as described above. Cells were fixed with 4% paraformaldehyde for 15 minutes, permeabilized, and then blocked with 0.5% Triton X-100 in phosphate-buffered saline containing 100 µg/mL of goat γ-globulin (Jackson Immunoresearch Laboratories) for 1 h at room temperature. Cells were incubated overnight at 4 °C with rat anti-HA antibody (1:500 - Roche) and then with secondary goat anti-rat IgG antibody coupled to Alexa 568 (Invitrogen/Life Technologies) for 1h at room temperature. Slides were examined by confocal microscopy using a confocal server spinning disc Zeiss platform equipped with a ×100 objective.

## Statistical analysis

The statistical analyses were performed with JMP pro 13 software. If more than two groups were compared, for parametric test ANOVA with post hoc tests as indicated were used or for non-parametric test Kruskall-Wallis with post hoc tests as indicated were used. (*, p < 0,05; **, p < 0,01; ***, p < 0,001). The number of biological replicate (n) analyzed is indicated in figure legends along with the number of independent experiment (N).

## Modeling dimers with different interfaces

Helical wheel diagrams have been drawn from grigoryanlab website (https://grigoryanlab.org/). Modeling of the cc-del6 and cc-del7 were performed with PyMol software and readapted from the structure 2loh.pdb (Nadezhdin et al., 2012).

## Acknowledgements

We thank Dr. Jean-Philippe Defour for initial lentiviral cloning of coiled coil dimeric proteins, Joanne Van Hees and Lidvine Genet for expert technical assistance and Dr. Nicolas Dauguet and Dr. Xavier Cahu for Flow Cytometry support. SNC is Honorary Research Director at FRS-FNRS Belgium. Funding to SNC is acknowledged from Ludwig Institute for Cancer Research, Fondation contre le cancer, Salus Sanguinis and projects Action de recherche concertée (ARC) 16/21-073 and WelBio F 44/8/5 - MCF/UIG – 10955. Funding to PKC is acknowledged from SAO-FRA Alzheimer Research Foundation. FP was supported by ARC 16/21-073 (to SNC) and ARC14/19-059 (to PKC). Support funds from FNRS grant PDRT.0177.18 to PKC, from NIH AG27317 to SOS and subcontract to NIH AG27317 to SNC are acknowledged.

## Author contributions

FP, NP^(a)^, RO and CV designed and performed experiments and interpreted data, PKC, SNC and SOS designed experiments, defined the strategy of the study, and interpreted data, FP, NP^(a)^, PKC, SOS and SNC wrote the paper, DV was responsible for mass spectrometry experiments, DMV and NP^(b)^ provided reagents and interpreted data. NP^(a)^: Nicolas Papadopoulos NP^(b)^: Nathalie Pierrot

## Supplementary Materials

**Supplementary Figure 1.**
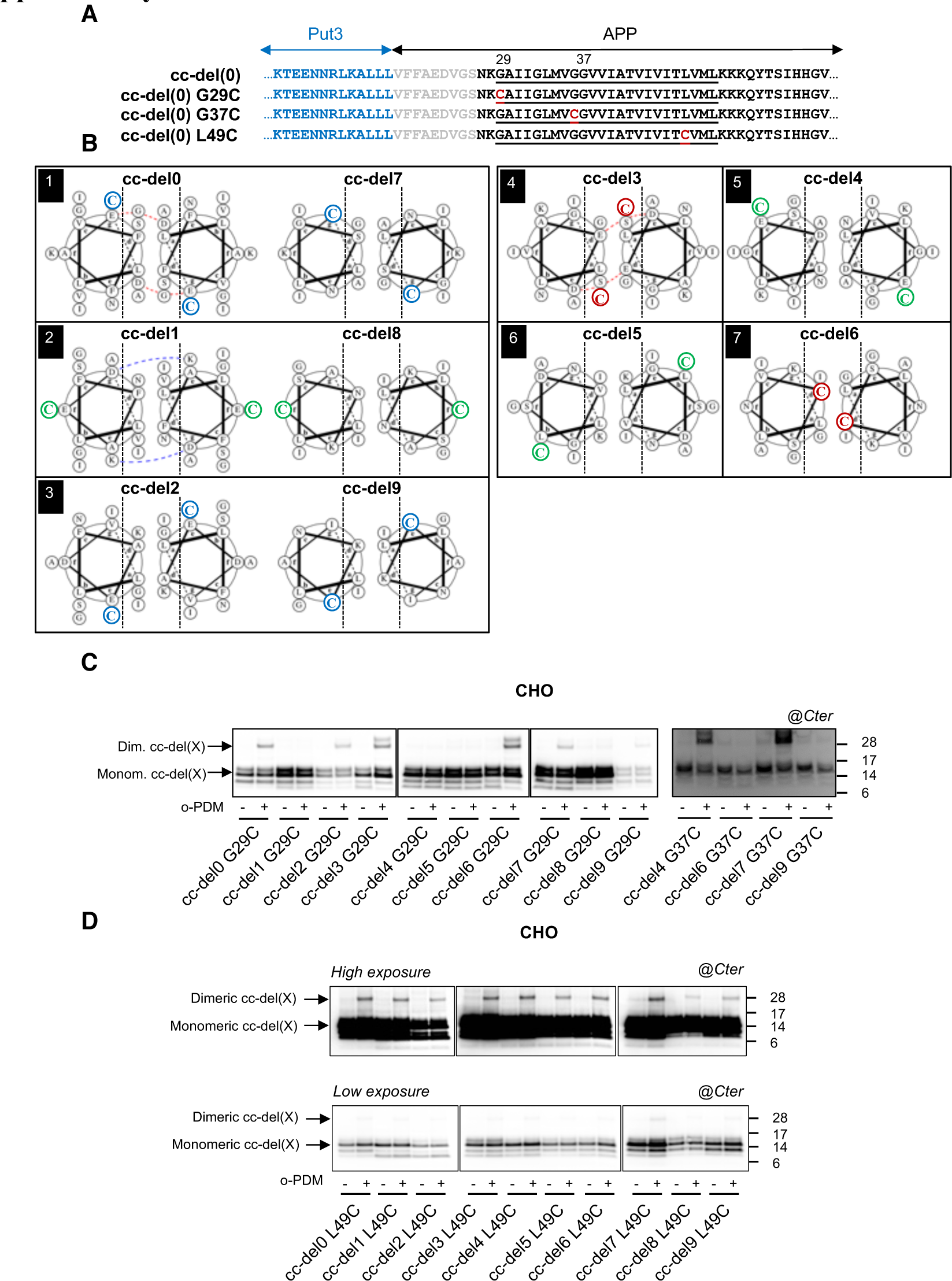
Validation of the coiled-coil C99 fusion proteins. **A.** Representation of the cc-del(X) and the corresponding cc-del(X) G29C, G37C and L49C sequences with cc-del0 as an example. Residues deleted in cc-del(X) constructs are in grey. Introduction of cysteine at position 29 (G29C) is highlighted in red. The mutation allows cross-linking with N,N’-1,2-phenylenedimaleimide (o-PDM) if two cysteines residues are close enough. **B.** Alpha helical prediction of the precise position of amino acids residues with the cysteine mutation at Gly29. Cross-linking predictions are indicated by a color code associated to the position Cys29 in the helical diagram. Cys29 is indicated in red, blue or green if it is present at the interface, at the border of the interface or outside the interface, respectively. **C.** Crosslinking study on living cells. CHO cells were transfected with the ten cc-del(X) G29C vectors or four cc-del(X) G37C vectors and treated (+) or not (-) o-PDM. Cell lysates were analyzed by Western Blotting and revealed with APP Cter antibody. **D.** Crosslinking study on living cells. CHO cells were transfected with the ten cc-del(X) L49C vectors and treated (+) or not (-) with o-PDM. Cell lysates were analyzed by Western Blotting and revealed with APP Cter antibody. Top: high exposure, bottom: low exposure.

**Supplementary Figure 2.**
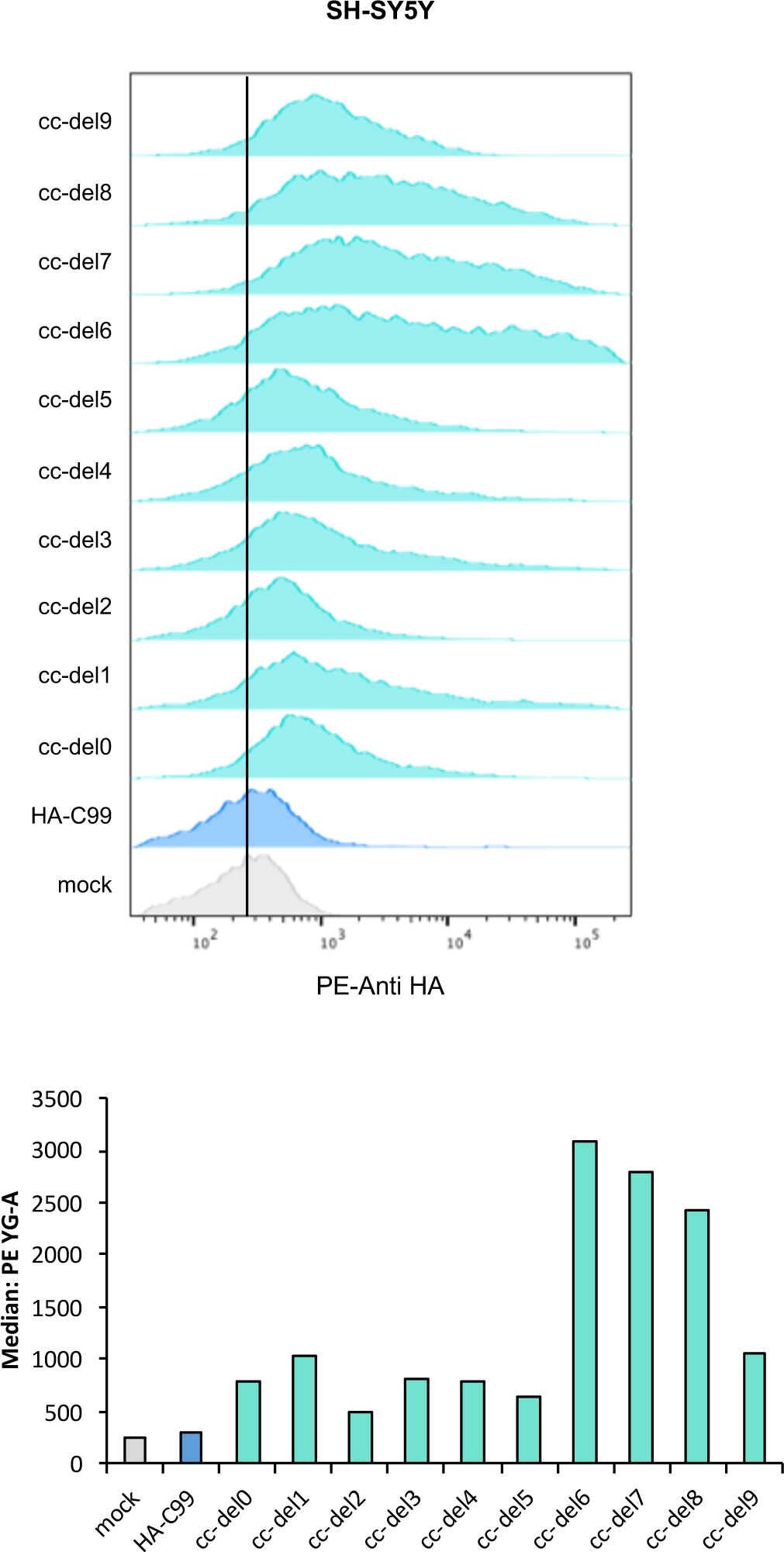
Surface expression of cc-del(X) constructs. FACS analysis of SH-SY5Y stably expressing HA-C99 and cc-del(X). Surface HA level of all SH-SY5Y was assessed by FACS using a PE-linked HA antibody. The histograms represent one representative experiment. The black line represents mean of the SH-SY5Y mock.

**Supplementary Figure 3.**
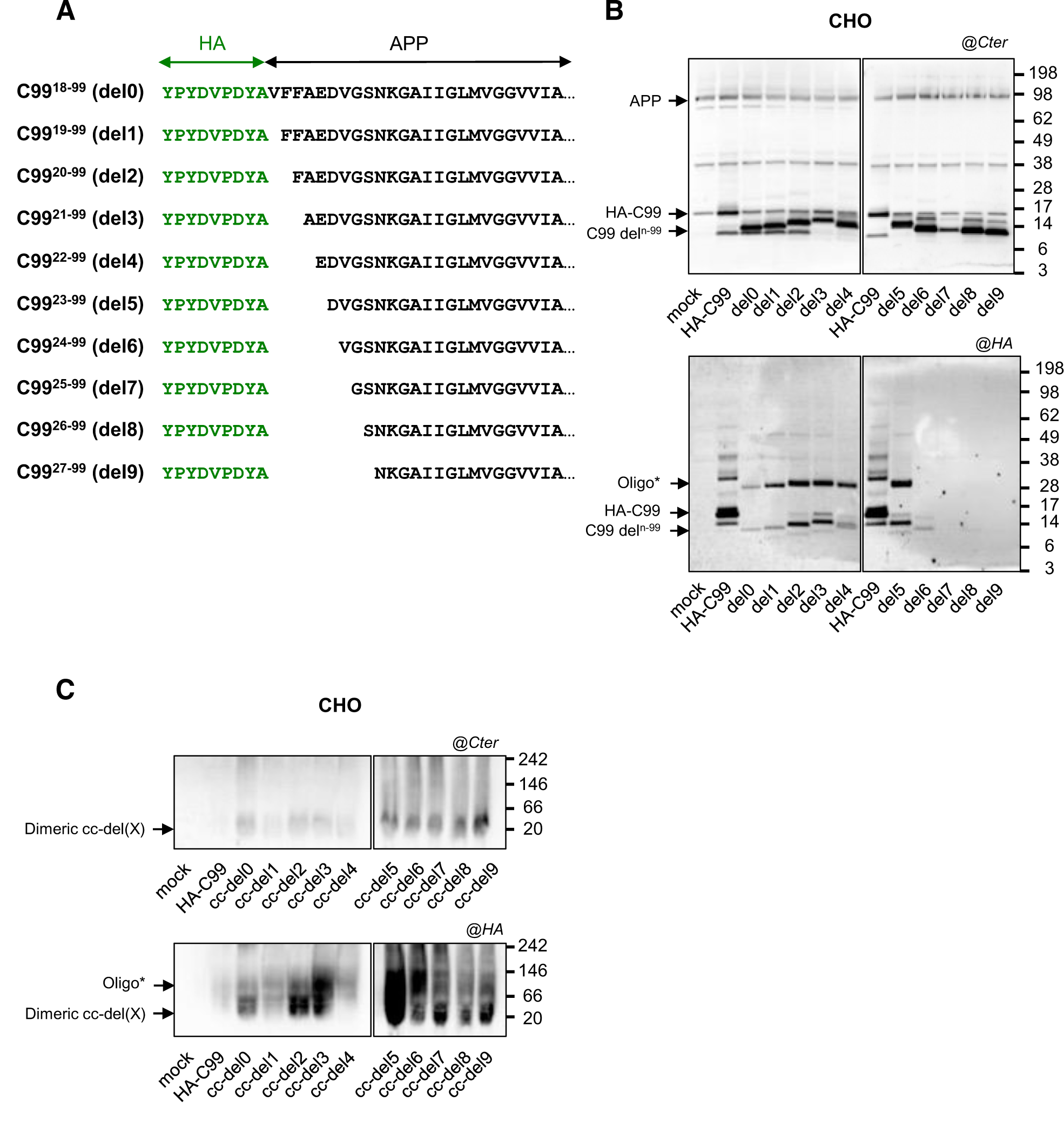
Oligomerization properties of C99^n-99^ and cc-C99^n-99^. **A.** Deletions of amino acids 18 to 27 in the C99 sequence followed by fusion with HA-tag in the N-terminal part. **B.** Profile of deletion in the N-terminal of C99. CHO cells were transiently transfected with mock vector, HA-C99 and one of the ten truncated C99 (where n is the first amino acid of the sequence). Cells were lysed and analyzed by SDS-PAGE followed by immunoblotting with Cter APP antibody (top) or HA antibody (bottom). **C.** Profile of coiled coil in native condition. CHO cells were transiently transfected with cDNA of mock vector, HA-C99 and the ten coiled coil constructs. Cells were lysed and analyzed on native page followed by immunoblotting with APP Cter antibody (top) or HA antibody (bottom).

**Supplementary Figure 4.**
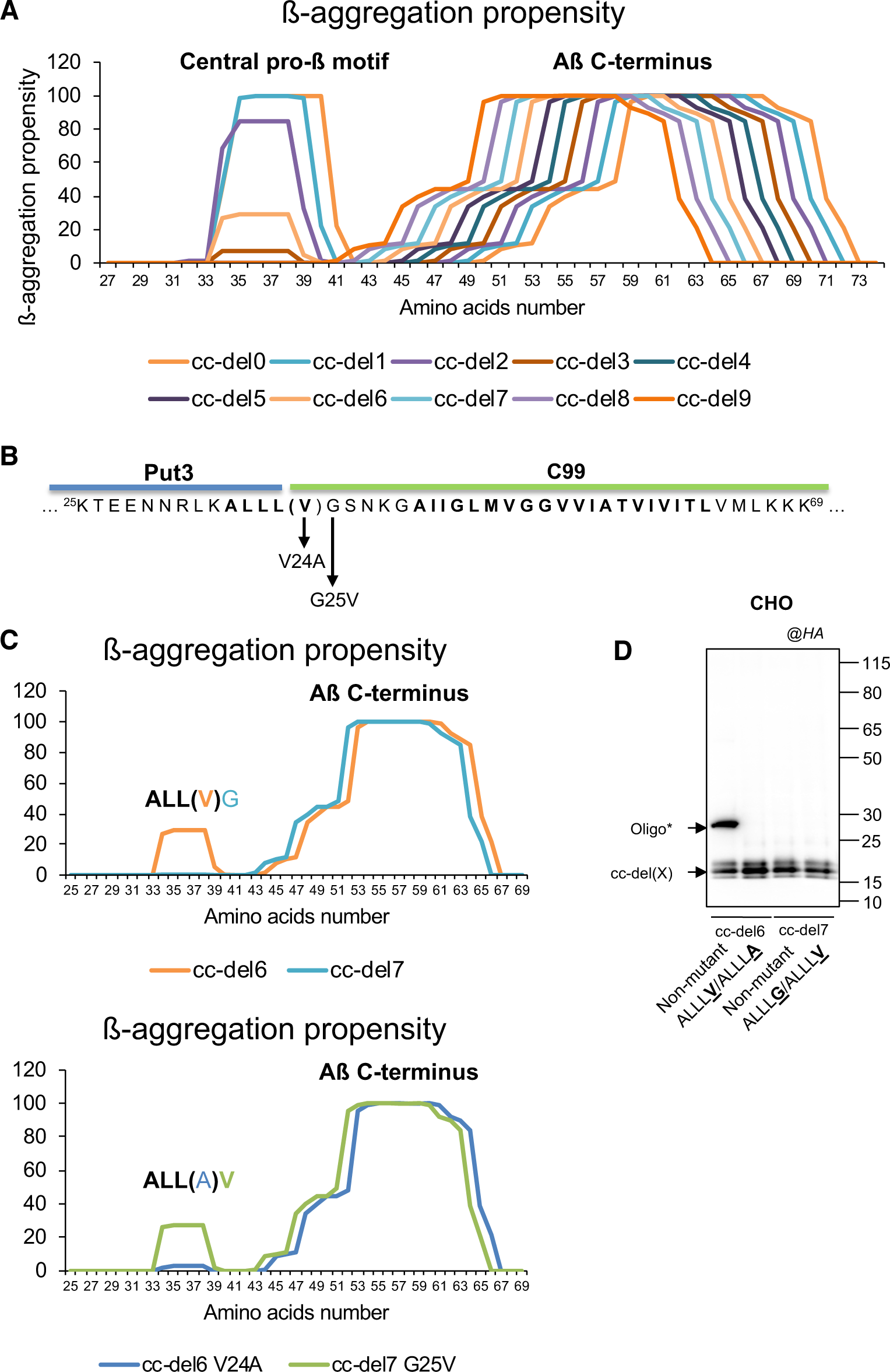
β-aggregation propensity of cc-del(X) constructs. **A.** Beta-aggregation propensity prediction using the Tango algorithm (Fernandez-Escamilla et al., 2004) for cc-del(X) constructs. **B.** Partial sequence of cc-del6/7 constructs. The additional amino acid (Val24) present in cc-del6 compared to cc-del7 is shown in bracket. Mutations are indicated by arrows. Amino acid sequences predicted to have a high propensity for cross-β aggregation are in bold. **C.** β-aggregation propensity prediction of cc-del6 and cc-del7 with and without the V24A and G25V mutations, respectively. Analysis done using the Tango algorithm (Fernandez-Escamilla et al., 2004). **D.** Analysis of the profile of cc-del(X) constructs and Aβ-like peptides in denaturing conditions. CHO cells were transiently transfected with mock vector, HA-C99, the cc-del6 and cc-del7 constructs with or without the indicated mutation or their respective truncated forms at position 42 or 43. Cell lysates were analyzed by Western Blotting and revealed with HA antibody.

**Supplementary Figure 5.**
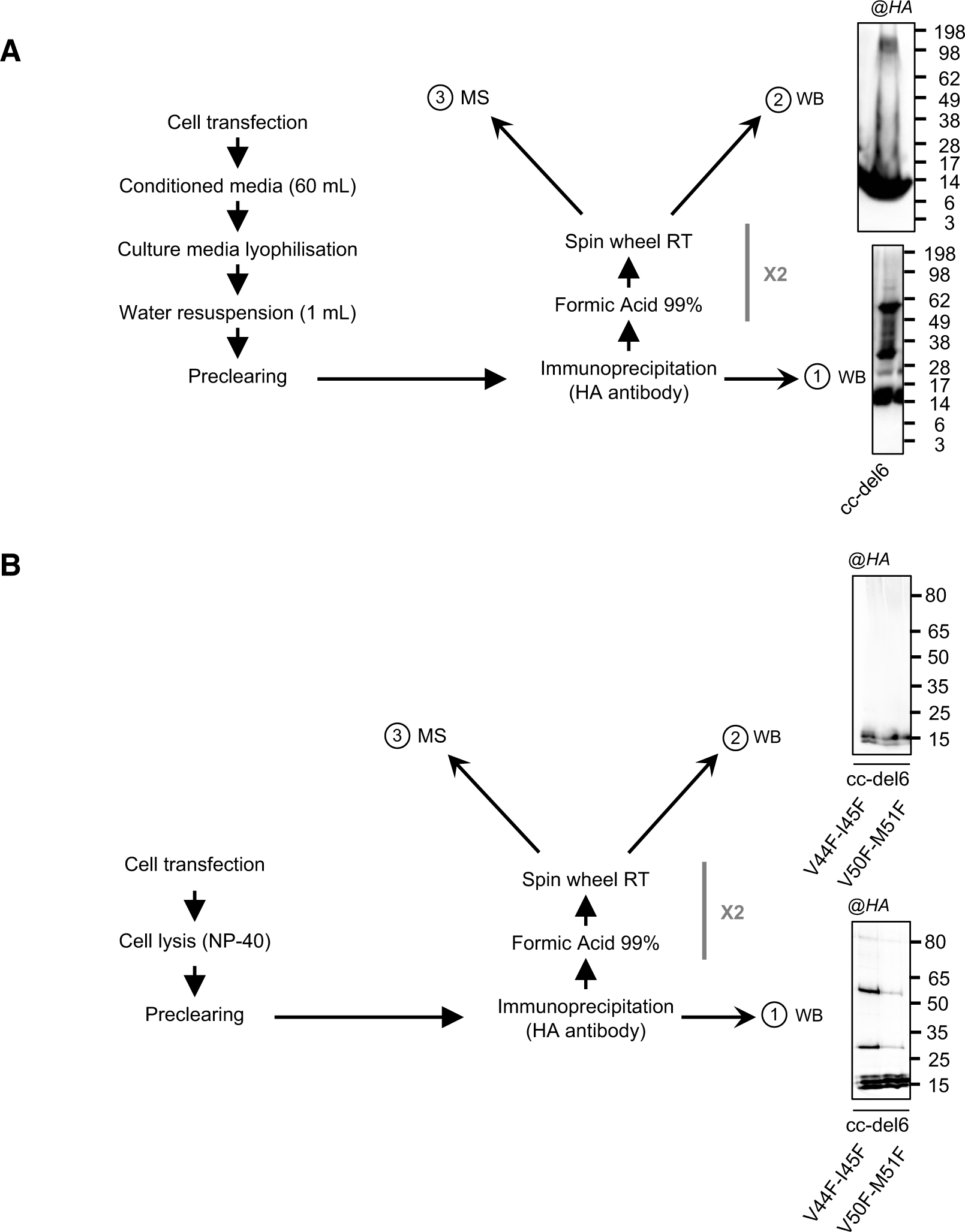
Sample preparation for mass spectrometry. **A.** Sample preparation for mass spectrometry (MS) analysis of oligomers associated with cc-del6. Conditioned media of CHO cells expressing cc-del6 were immunoprecipitated and analyzed by Western blotting before (1) or after treatment with formic acid (2). Samples treated with formic acid were further analyzed by tandem mass spectrometry (MS/MS). **B.** Sample preparation for mass spectrometry (MS) analysis for oligomers associated for cc-del6 V44F-I45F and cc-del6 V50F-M51F. Cell lysates of CHO cells expressing cc-del6 were immunoprecipitated as described in (A) and analyzed by Western blotting before (1) or after treatment with formic acid (2). Samples treated with formic acid were further analyzed by tandem mass spectrometry (MS/MS) (3). For each condition, the lysates of three confluent 15cm^2^ dishes were pooled.

**Supplementary Figure 6.**
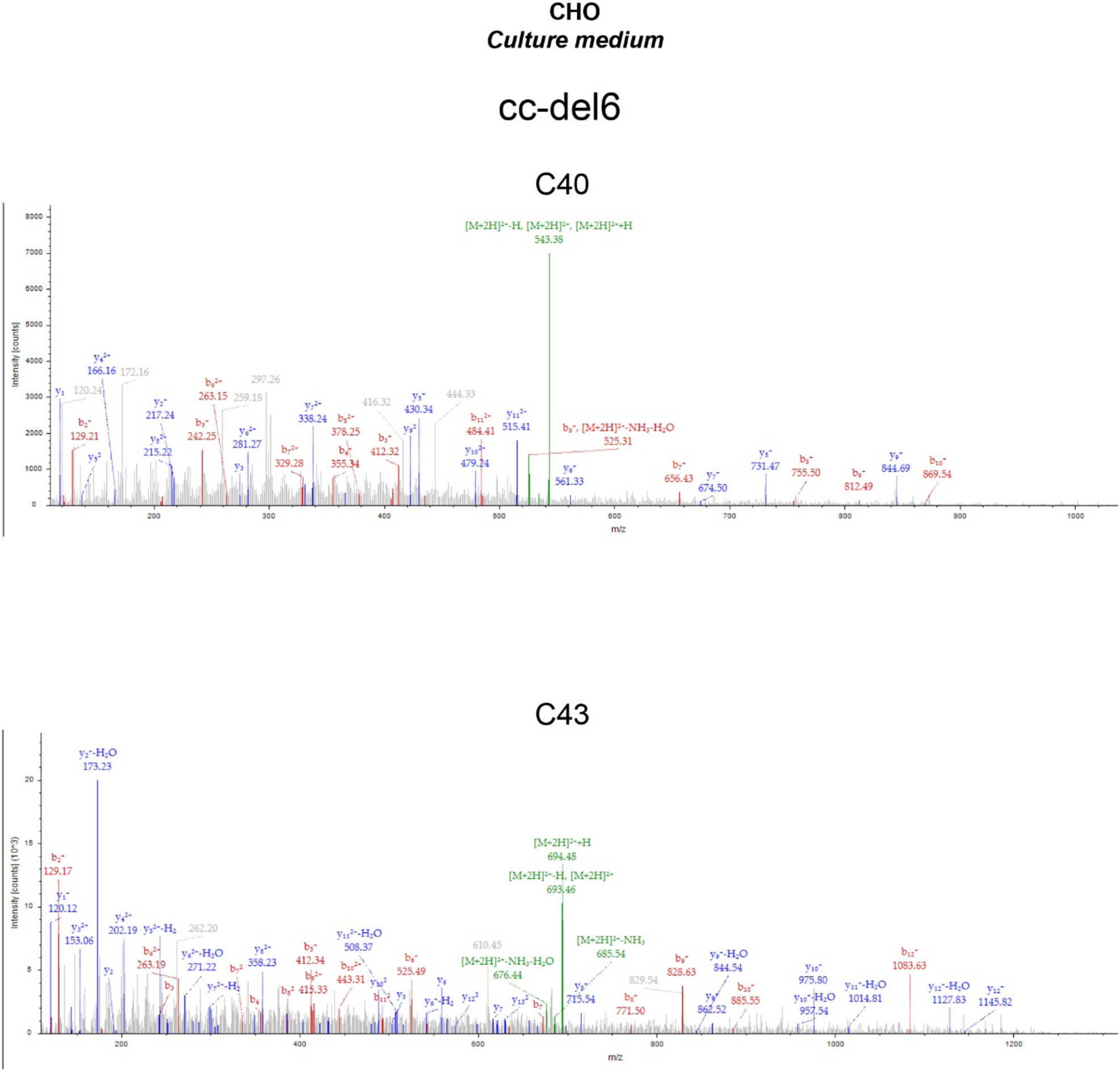
Identification of Aβ C-terminus in cc-del6 by LC-MS/MS. Identification of C40 and C43 peptides by LC-MS/MS. The LC-MS spectrum shows MS^2^ obtained from a +2 charged parent ion of m/z 543.3223 (C40) and 693.902 (C43) respectively, which upon HCD fragmentation gave rise to the y- and b-series of daughter ions allowing confirmation of the identified peptide sequences.

**Supplementary Figure 7.**
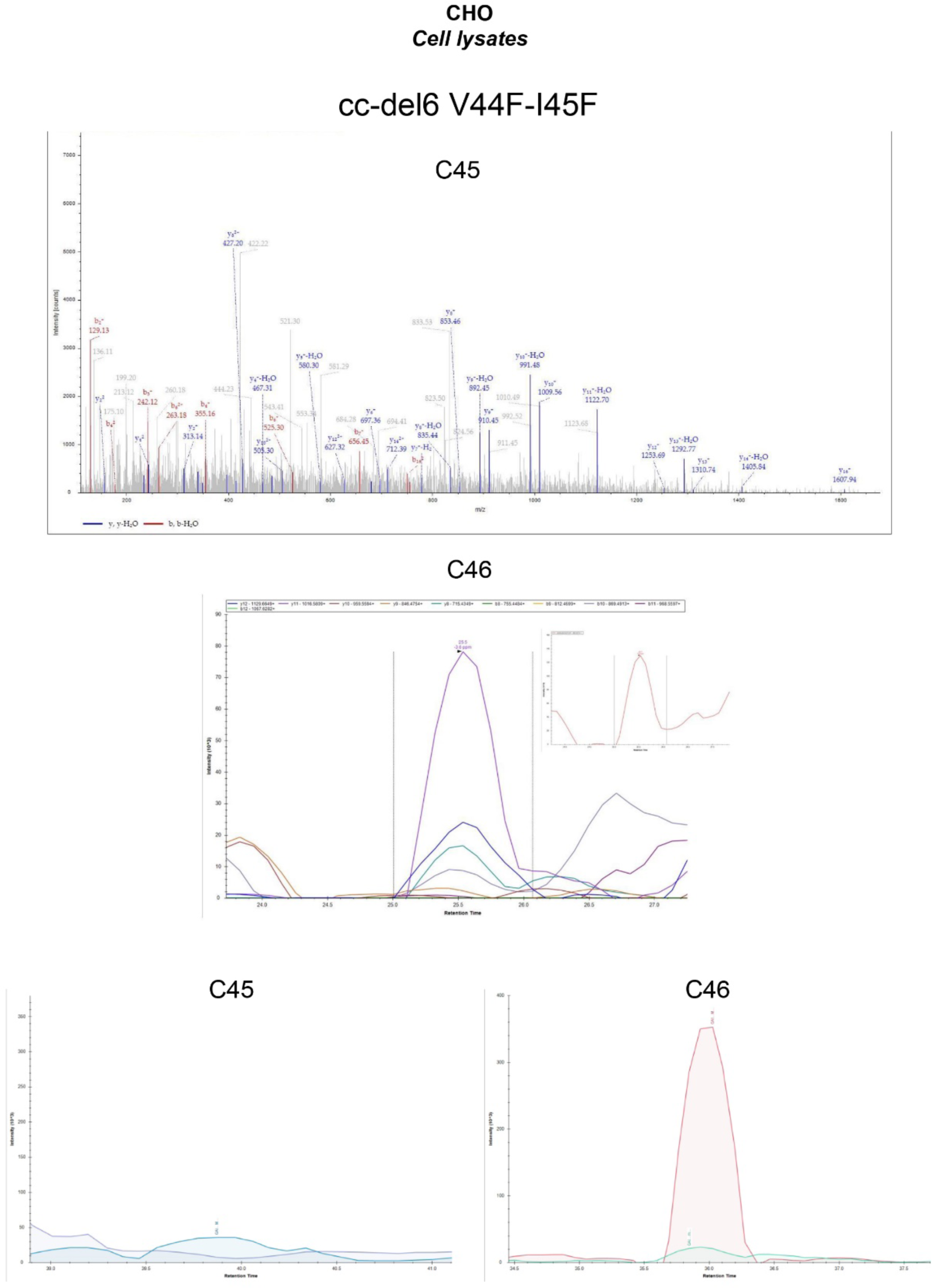
Identification of Aβ C-terminus in cc-del6 V44F-I45F by LC-MS/MS. **A.** Identification of C45 and C46 peptides by LC-MS/MS. The upper LC-MS/MS spectrum shows MS^2^ obtained from a +2 charged parent ion of m/z 832.982 (C45), which upon HCD fragmentation gave rise to the y- and b-series of daughter ions allowing confirmation of the identified peptide sequences. The lower chromatogram shows peak area contribution from a +2 precursor charged parent ion of m/z 882.5069 (C46) (small insert) and the corresponding coeluting transitions for 10 y- and b-daughter ions determined by a PRM experiment. **B.** Relative quantification of C45 and C46 peptides. The panels show the precursor ions intensities from C45 and C46 parent ions determined from the PRM experiment

**Supplementary Figure 8.**
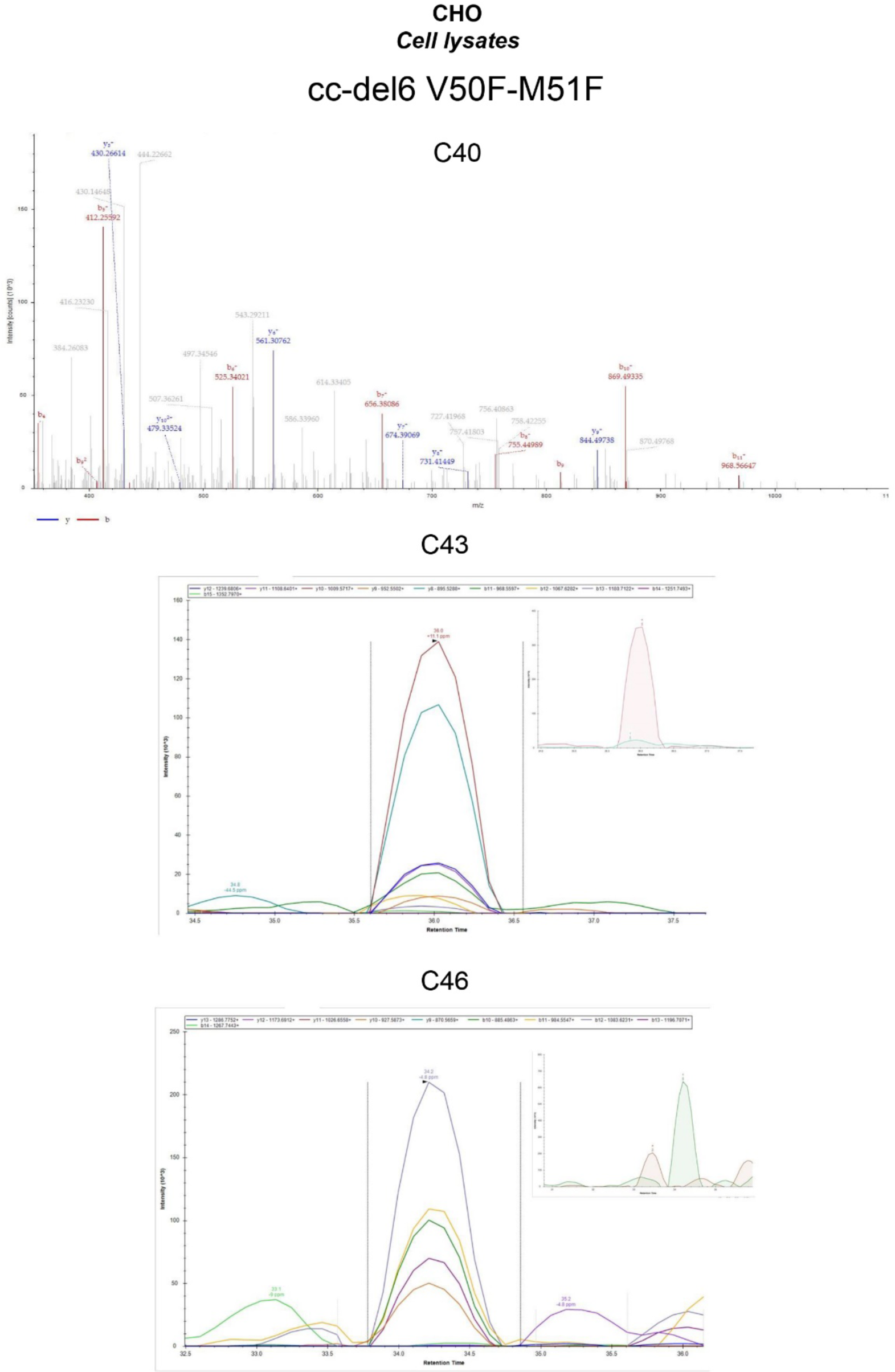
Identification of Aβ C-terminus in cc-del6 V50F-M51F by LC-MS/MS. Identification of C40, C43 and C46 peptides by LC-MS/MS. The upper LC-MS/MS spectrum shows MS^2^ obtained from a +2 charged parent ion of m/z 543.3228 (C40), which upon HCD fragmentation gave rise to the y- and b-series of daughter ions allowing confirmation of the identified peptide sequences. The lower chromatograms show peak area contribution from a +2 precursor charged parent ion of m/z 693.9118 (C43) or 847.5111 (C46) (small inserts) and the corresponding coeluting transitions for y- and b-daughter ions determined by a PRM experiment.

**Supplementary Figure 9.**
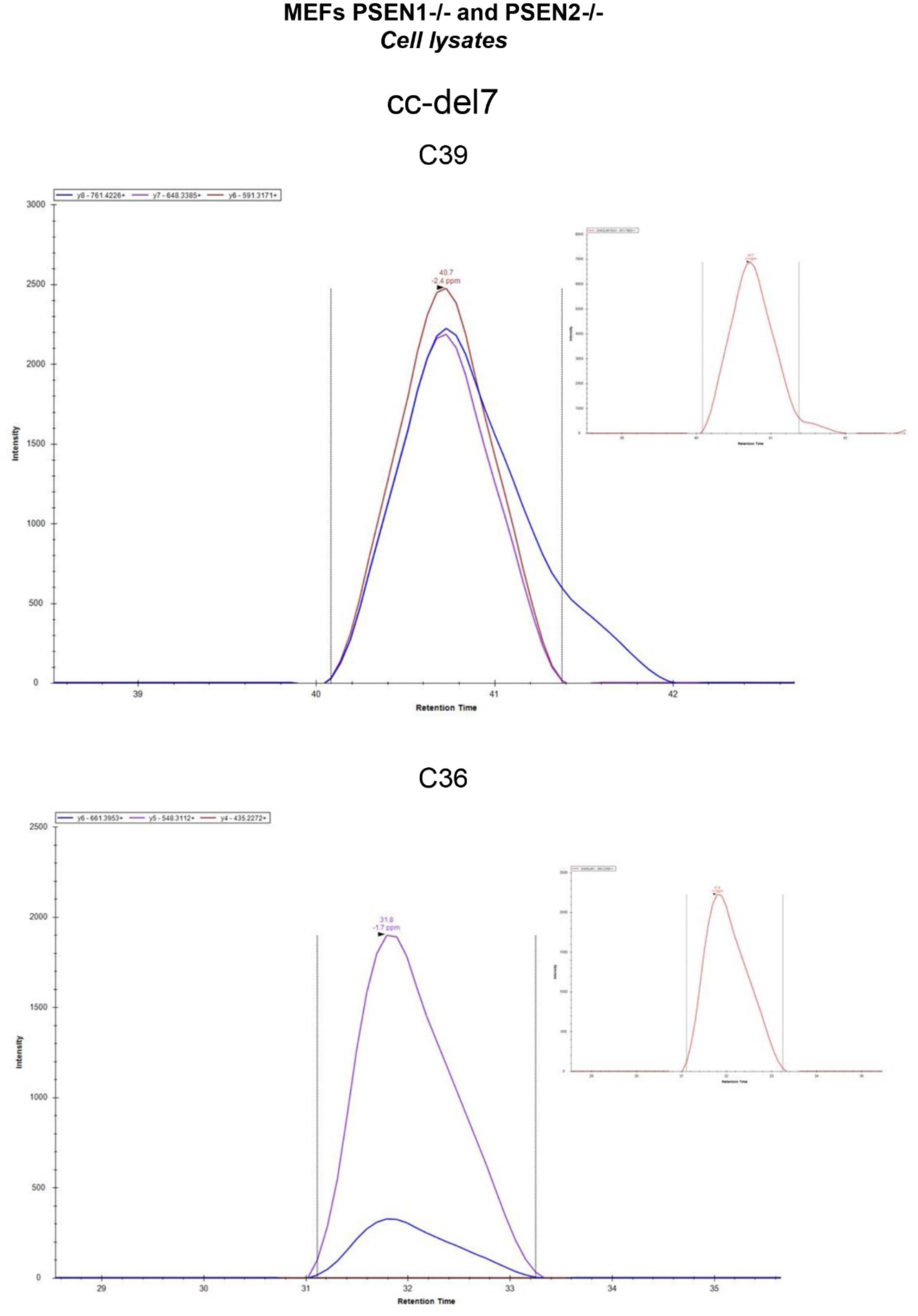
Identification of Aβ C-terminus in cc-del7 by LC-MS/MS. Identification of C39 and C36 peptides by LC-MS/MS.. The chromatogram shows peak area contribution from a +2 precursor charged parent ion of m/z 493.7875 (C39) or m/z 387.2322 (C36) (small inserts) and the corresponding coeluting transitions for y- and b-daughter ions determined by a PRM experiment

